# Directed growth and fusion of membrane-wall microdomains requires CASP-mediated inhibition and displacement of secretory foci

**DOI:** 10.1101/2022.07.03.498613

**Authors:** ICR Barbosa, D De Bellis, I Flückiger, E Bellani, M Grangé-Guerment, K Hématy, N Geldner

## Abstract

Casparian strips (CS), the main extracellular diffusion barrier in plant roots, are precisely localized cell wall lignin-impregnations, contrasting animal tight-junctions. The CS membrane domain (CSD) proteins 1-5 (CASP1-5) define and accumulate at the CS associated membrane domains displaying matrix adhesion and protein exclusion. A full CASP knock-out (*caspQ*) now reveals that CASPs are not needed for localization of lignification or lignin-polymerizing enzymes, since correctly aligned spots still form in the mutant. Ultra-structurally, however, these spots appear as highly disorganized secretory foci, with neither exclusion zone nor membrane attachment and excessive cell wall growth. Biotin proximity labelling identifies RabA-GTPases as potential CASP-interactors. We confirm their localisation and function at the CSD, similar to exocyst subunits, known Rab effectors. Our work reveals that CASPs enforce displacement of initial secretory foci through exclusion of vesicle tethering factors, thereby ensuring rapid fusion of microdomains and effective sealing of the cell wall space.

## Introduction

Plants acquired multicellularity independently of animals and thus evolved independent solutions to the many challenges that arose with multicellularity (Knoll, 2011). One fundamental feature of plants is the presence of a cell wall that allows plant cells to resist high internal pressure, enabling rapid cell growth through a dominating central vacuole (Somerville et al., 2004). Yet, cell walls necessarily immobilise and isolate cells, not allowing for cell-to-cell contact of plasma membranes, something which animals rely on extensively during development and cellular signalling. Thus, many of the profound differences between animal and plant development can be seen as resulting from the fact that plants have generated complex multicellularity from walled cells. A particularly straightforward difference arising from this, is the way polarised epithelia can be formed. Polarised epithelia restrict extracellular diffusion between different extracellular spaces by generating specific, extracellular diffusion barriers. As a consequence, epithelia have to maintain two functionally distinct plasma membrane surfaces, facing the different extracellular environments, allowing them to generate selective and vectorial transport. In animals, extracellular diffusion barriers are formed by tight and adherence junctions, polymeric assemblies of transmembrane proteins that enter in direct contact between neighbouring epithelial cells, forming a network of joined rings (Cereijido, 2004).

Because of their cell walls, plants cannot achieve extracellular (apoplastic) diffusion barriers by membrane protein-mediated cell-to-cell contacts. Instead, a more complex interplay between transmembrane proteins and cell wall enzymes must generate a localised, coordinated membrane-wall adhesion, guiding hydrophobic cell wall modifications between cells, thus creating a membrane-to-cell-wall-to-membrane continuum. We are only beginning to understand the complex molecular identity of this plasma membrane-cell wall nexus (Barbosa et al., 2019). The CS is an illustrative example of such a tight membrane-wall barrier. Work from our lab initially established that the CS is made of lignin - a hydrophobic, polyphenolic polymer, impregnating cellulosic plant cell walls (Naseer et al., 2012). We then discovered the CASPARIAN STRIP MEMBRANE DOMAIN PROTEINs (CASPs) - the first proteins localised to the CS – and showed that their function is to form an extended, highly stable transmembrane platform (Roppolo et al., 2011). CASPs form central, longitudinal rings in each endodermal cell, with their coordinated localisation leading to a net-like appearance at a tissue-wide level that precisely coincides with the lignin impregnations of the CS themselves. These CASP platforms are thought to bring together reactive oxygen species (ROS)-producing NADPH oxidases with ROS-utilising peroxidases, both of which have been shown to be crucial for CS formation (Lee et al., 2013; Rojas-Murcia et al., 2020). A number of other cell wall-localized proteins, such as ESB1 have also been identified to both localize to the CS and to be involved in its formation (Hosmani et al., 2013; Reyt et al., 2020), indicating that a complex machinery is involved in locally synthesising CS lignin. CASPs are small, four-transmembrane-span proteins, whose features, such as very strong endodermis-specific expression, high stability, lack of lateral mobility, strong association with cell wall containing fractions, as well as extensive cross-interactions in *in vivo* pull downs, all pointed to a central, structural role of these proteins in the formation of the CSD (Roppolo et al., 2011). However, evidence for physical interactions between CASPs and cell wall enzymes has remained indirect. ESB1, for example, a cell wall protein carrying a dirigent-like domain, is required for CASP stabilisation, with the mutant displaying unstable CASP domains that do not fuse (Hosmani et al., 2013). However, no evidence for direct interaction of ESB1 and CASPs has been reported. Currently, only partial loss-of-functions for the CASP gene family have been reported. A *casp1 casp3* double mutant displayed a clearly interrupted CS and strong, ectopic lignification of cell corners (Lee et al., 2013; Roppolo et al., 2011). Enhanced ectopic lignification is now understood not to be a direct result of reduced CASP function. Rather, it is due to the activation of a signal transduction pathway (the SCHENGEN (SGN) pathway), whose role is to surveil the integrity of the extracellular diffusion barrier, to boost Casparian strip formation and to initiate compensatory lignification in case of defects (Alassimone et al., 2016; Doblas et al., 2017; Fujita et al., 2020; Pfister et al., 2014). Activation of the SCHENGEN pathway in the double mutant clearly indicates that reduction-of-function of CASPs causes barrier defects. However, it was unknown whether the remaining, centrally aligned lignin foci are due to a CASP-independent localisation and activity of lignin enzymes or whether they simply reflected the activity of the remaining CASPs in this mutant. Indeed, knock-out of the transcription factor MYB36 prevents expression of all CASPs (Kamiya et al., 2015). This is associated with a complete absence of CS and their associated membrane domain. Yet, lack of MYB36 activity affects expression of more than hundred genes, some of which are known to be involved in lignification, CASP stability or signal transduction, suggesting that MYB36 controls large parts of the endodermal differentiation program and that its phenotype cannot be subsumed to a specific absence of CASP function.

Finally, the strongest CASP1 delocalisation phenotype was observed in a mutant of an exocyst subunit (*exo70a1/lotr2*) (Kalmbach et al., 2017). The exocyst is a conserved eukaryotic complex that is required for tethering late secretory vesicles to the plasma membrane, enabling their fusion. In yeast and animal systems, it has been demonstrated that Rab GTPases mediate vesicle fusion through recruitment or activation of exocyst subunits to secretory vesicles. Plants have undergone a specific, large irradiation of EXO70 subunits, with 23 homologs in Arabidopsis (Žárský et al., 2009). This has led to the speculation that in plants, different EXO70 subunits are able to direct specific subsets of vesicles to distinct membrane domains. Indeed, in *exo70a1*, CASPs accumulate in small, unstable microdomains all over the endodermal surface (Kalmbach et al., 2017). This indicates that EXO70A1 is crucial for localisation, yet not for CASP1 secretion *per se*, which could occur through other EXO70 subunits. Intriguingly, a tagged version of EXO70A1 transiently accumulate at the site of CASP microdomain formation, slightly preceding CASP1 localisation itself. This suggested a model whereby, in the endodermis, EXO70A1 mediates the selective, localized tethering of CASP-bearing vesicles to the median plasma membrane domain.

Our in-depth analysis of a full knock-out of CASP activities, together with our identification of RabA proteins as new players in CS formation, now allows us to draw a very different model of CASP action, in which CASPs are not needed for localisation or activity of lignification enzymes *per se*. Rather, CASPs are needed to form a membrane domain of protein exclusion and cell wall attachment that rapidly suppresses further secretion to the same initial focus by evicting EXO70A1 secretion landmarks and forces displacement of secretory foci along the median line, thus ensuring rapid fusion of foci into an uninterrupted band. Moreover, CASP action also appears to be required to ensure organized and spatially limited lignification, allowing for the formation of the idiotypic, flat lignin structure of specific width that motivated the use of the term “strip” in its original description.

## Results

### Endodermal knockout of CASP function leads to abnormal lignin foci

To identify CASP function in Casparian Strip formation we aimed to isolate a full endodermal *CASP* mutant. CASPs belong to the family CASP-LIKES, with 39 members in Arabidopsis, belonging to the super family of eukaryotic MARVEL proteins (Roppolo et al., 2014). CASP1 – CASP5 were categorized based on their strong, endodermis-specific expression and localization to the CSD. Since there were no available T-DNA, or other mutants, for all CASP genes, we isolated a double mutant, based on two strong, exon-localized T-DNA insertion alleles, *casp3-1 casp5-1* and targeted the remaining three CASPs by multiplexing CRISPR-Cas9 (Supplementary Figure 1.1) (Ursache et al., 2021a). With two gRNAs targeting each CASP1, 2 and 4, we retrieved a *casp quintuple* (*caspQ*) mutant, heteroallelic for the three genes, deletion and frame-shift mutation alleles, from which we isolated three homozygous allelic combinations (Supplementary Figure 1.1). Firstly, we measured extracellular (apoplastic) barrier function by measuring penetration of Propidium Iodide (PI) into the central vasculature (Figure 1a). Using this established diffusion barrier assay, we found *caspQ* plants had a delay in the establishment of the apoplastic barrier within the typical range for mutants with interrupted (*casp1-1 casp3-1* and *esb1-1*) or absent CS (*myb36*). This is due to a compensatory mechanism, dependent on the activation of the SGN3 (GSO1) receptor kinase, which leads to the formation of ectopic lignification and delayed sealing of the apoplastic space (Pfister et al., 2014). Consequently, *sgn3* mutants have a virtually absent apoplastic barrier throughout the whole root and were used as a control (Figure 1a). Next, we used a fluorescent lignin stain (Basic Fuchsin) to highlight the CS structure of *caspQ* (Figure 1b). Surprisingly, with this assay, the novel *caspQ* displayed a similar phenotype to the previously described *casp1 casp3* double mutant, i.e. interrupted lignin foci and the typical compensatory ectopic cell corner lignification (Barbosa et al., 2019; Roppolo et al., 2011). This impression was confirmed by quantitative image analysis (1c-e). At 20 cells after onset of elongation, wild-type (Col-0) CS displayed one particle per region of interest, indicating complete fusion at this stage. As images suggested, we found statistic differences with increasing complexity of *casp* mutants. We found that double *casp1 casp3,* the triple *casp1 casp3 casp5,* and *esb1* displayed the highest number of particles, while *caspQ* showed a lower number of particles, but occupied a similarly reduced total area. Thus, *caspQ* has less, but bigger particles than lower order mutants (Figure 1b-c). In wild-type, the forming CS is deposited initially in centrally aligned dots, coined “string-of-pearl” stage, that rapidly progress into a fused strip. To understand whether the number of dots in *casp* mutants reflected a halted progression of CS development, we conducted a developmental progression analysis in selected genotypes (Col-0, *casp1 casp3 casp5* and *caspQ)*, analysing cell 9 (start of detectable Basic Fuchsin signal), 10, 15 and 20 (Figure 1d-e). As expected, wild-type displayed a quick progression in particle size from 7% at cell 9, to 100% at cell 20 (indicating complete fusion into one contiguous band). In contrast, both triple *casp1 casp3 casp5* and *caspQ* mutants failed to progress in CS coverage with average particles size remaining at around 3-6% size and a total coverage of about 50% relative to wild-type at cell 20, despite strong variation (Figure 1e). This analysis supports the idea that CS development in *casp* mutants is arrested at the initial string-of-pearl stage.

**Figure 1:**
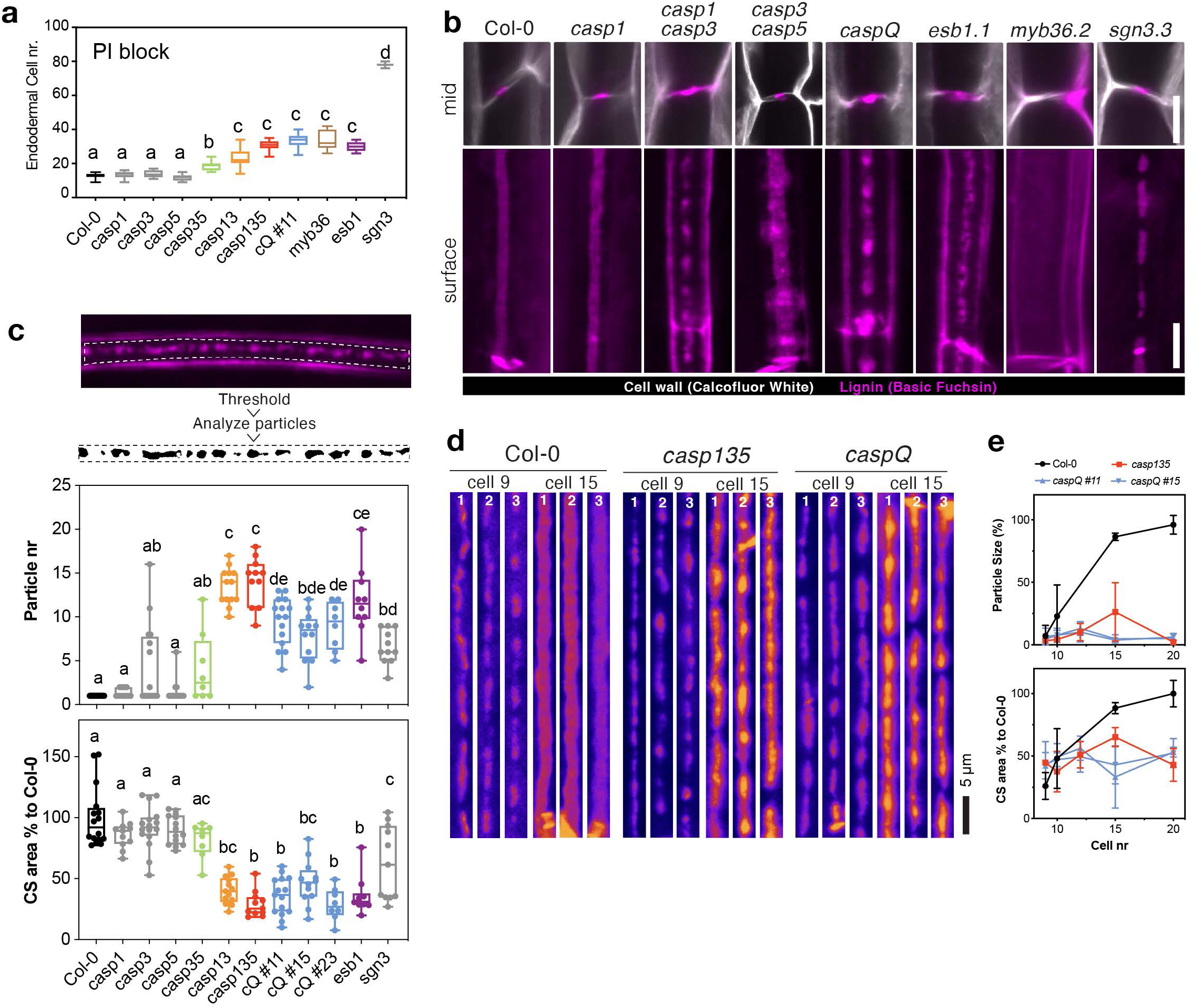
Casparian Strip development arrests at initiation sites in caspQ. **a.** Propidium iodide uptake assay, endodermal cell number indicates position at which uptake is blocked. Mean, standard deviation (n=10) and significant differences to wild-type Col-0 shown with different letter by ANOVA, Tukey test *p*-value < 0.05. **b.** Mid (upper) and surface (lower panel) views of CS-lignin (magenta, Basic Fuchsin) and cell wall (gray, Calcofluor White) staining in wild-type, CS mutants and *caspQ* at endodermal cell 20. **c.** Particle analysis in CS surface of endodermal cell 20, as described in Materials and Methods, with variables Particle number and CS % area to Col-0 measured per 46 µm of CS surface. Mean, standard deviation (8 ≤ n ≤ 15) and significant differences to wild-type shown with different letter by ANOVA, Tukey test *p*-value < 0.05. **d.** Representative pictures of CS surface views from three individuals at endodermal cell number 9 and 15 from Col-0, *casp135* and *caspQ*. **e.** Particle size (%) and CS area (%) relative to mean Col-0 at cell 20 was estimated as in c. Scale bars 5 µm.

These phenotypes of the *caspQ* mutants forced us to re-consider our long-standing notion that the remaining, centrally-aligned lignin foci observed in *casp* multiple mutants are due to the remaining activity of CASP homologs. Yet, since CASP1-5 belong to the multi-gene CASP-LIKE family of 39 members, we wanted a ascertain that none of these CASPLs could somehow compensate for the loss of CASPs in *caspQ* endodermis - either by normal, basal expression or through induced expression in the mutant - and thus be responsible for the remaining, centrally aligned lignin dots. For this, we inspected expression of the CASPL-family in several RNA-Seq datasets for endodermis-specific expression, and in mutants or treatments mimicking CS defect. These included *(i)* protoplast-sorted RNA-Seq data in wild-type and *myb36* (“myb36”) (Liberman et al., 2015), (*ii)* CIF2 treatment RNA-Seq (Fujita et al., 2020), (*iii)* targeted purification of polysomal mRNA with endodermal specific promoters (“endodermal TRAP”) (Andersen et al., *unpublished*). We found that seven other members of the CASPL1 clade, to which CASP1-5 belong, namely CASPL1A1, 1B1, 1B2, 1C1, 1C2, 1D1, 1D2, were either detected in endodermis-specific datasets and/or up-regulated in *myb36* mutant/conditions mimicking defective CS (CIF2 treatment) (Supplementary Figure 1.3). Indeed, some of these members were already reported to be expressed in suberizing endodermal cells (Champeyroux et al., 2019). However, expression analysis, using fluorescent CASPL fusion proteins in both wild-type and *caspQ* mutant background revealed: either no endodermal expression (1D1) or an onset of expression too late to be responsible for the initiation of the lignin dots seen in *caspQ* mutants (Supplementary Figure 1.3). Yet, to completely rule out the contribution of these genes for CS formation, we knocked-out the six endodermis-expressed members, (CASPL1A1, 1B1, 1B2, 1C1, 1C2, 1D2) in the *caspQ* generating the *undecuple* mutant, *caspQ 6x-caspl*. Yet, even in this mutant, we could not observe any increased phenotypic severity compared to *caspQ* (Supplementary Figure 1.3). We therefore concluded that *caspQ* is indeed a full *CASP* knockout and that localization and initiation of lignin deposition is independent of CASP activity.

### The centrally aligned dots in caspQ are hotspots of secretion and lignification

Next, we inspected the ultra-structure of *caspQ* dots by transmission electron microscopy (TEM) coupled with KMnO_4_ as a lignin stain. Here, we found that *caspQ* mutants display dramatic phenotypes in cell wall structure (Figure 2a). At 1.8mm from the root tip, wild-type CS appears as a thin lignified wall, sandwiched between the two tightly attached plasma membranes of neighbouring cells. *caspQ* mutants, by contrast, initially displays a spectrum of defects, from strong plasma membrane detachment with extracellular vesicles and irregular lignification (Figure 2a, root 1) to irregular, lignified cell wall thickenings and less pronounced, but still observable membrane detachment (Figure 2a, root 2). We found similar, but less frequent, membrane detachment and extracellular vesicles already in the triple *casp1 casp3 casp5,* but not in double *casp3 casp5* (Supplementary Figure 2.1a). Moreover, *esb1* also displayed an intermediate phenotype with significantly thickened, lignified cell wall and enhanced membrane detachment (Supplementary Figure 2.1a). By observing sequential developmental sections of the same seedling in wild-type (Col-0) and *caspQ* (Figure 2d-e), we found that the variability of the *caspQ* phenotype was due to a developmental progression. In wild-type, the CS forms approximately at 1mm from the root tip in 5-day-old seedlings. By capturing first images at 0.9 mm from the root tip in wild-type, we observed either of two stages: no difference in membrane or cell wall (stage 0), or invaginations and presence of extracellular, vesiculo-tubular bodies (EVBs), but with no obvious cell wall thickenings yet (stage 1) at the predicted position of the CS. In follow-up sections (1.1 mm), some cells displayed a thin CS, flanked by invaginations and EVBs on the edges (stage 2). Finally, at 1.2mm, we detected a wider, mature CS, with little to no EVBs on the flanks (stage 3) (Figure 2d,e). To our knowledge, such an ultra-structural analysis of progressive CS development has not yet been reported, especially the association of EVBs with early stages of CS formation. In *caspQ,* we found early stage (stages 0 and 1) to resemble wild-type. In the following sections, however, a modified wall became apparent that displayed thickening in all directions (as opposed to the strictly confined, thin strip of the wild-type). Associated with this was a continued presence of EVBs and strong membrane detachment. We categorized this as mutant Stage 2 (stage 2^m^). Only in later sections, around 2.4mm, EVB presence and membrane detachment finally abated with a strongly thickened, variably shaped cell wall at the position of the CS (Figure 2b,d). We categorized this as mutant stage 3 (stage 3^m^), as the final stage of central cell wall formation in *caspQ* (Figure 2d,e). In summary, this analysis revealed that *caspQ* appears to have a prolonged secretion phase, an inability of the plasma membrane to attach to the lignified wall and an uncontrolled cell wall outgrowth at the presumptive CS (Figure 2c,d).

**Figure 2:**
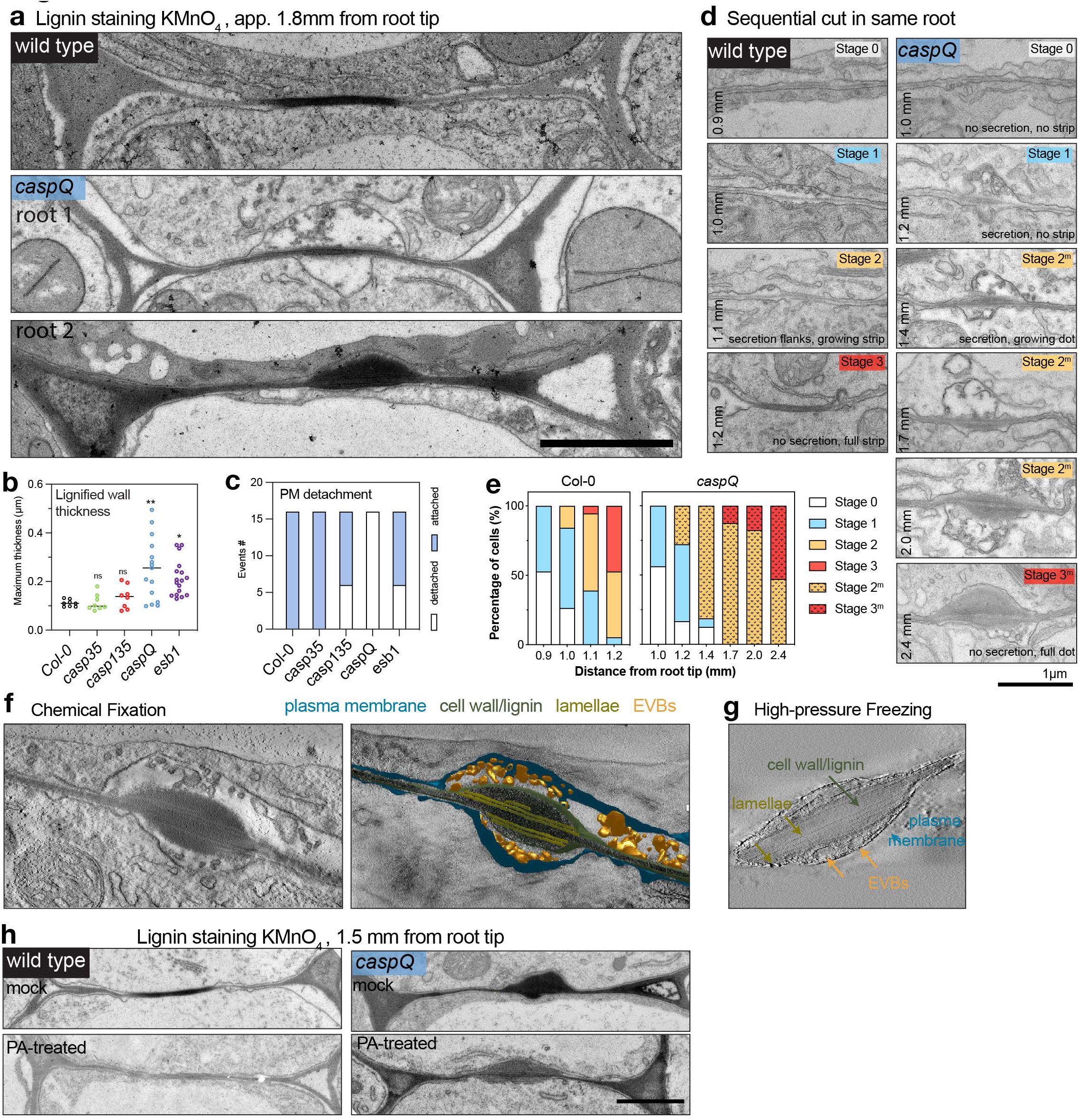
caspQ foci are lignin-rich secretion hotspots. **a-c.** Representative electron micrographs with KMnO_4_ lignin staining from wild-type and *caspQ* endodermal cell-cell junctions in transversal cuts at 1.8 mm from root tip. For *caspQ* two distinctive appearances are shown **(a)**. Quantification of maximum thickness of lignified cell-wall stained by KMnO_4_, mean (line, n=16) and significant differences to wild-type by ANOVA, Dunnett’s test *p*-value < 0.05. **(b)**. Total number of events of plasma membrane detachment in total 16 cells from two individuals **(c)**. **d-e.** Developmental staging of wild-type CS and *caspQ* foci. Pictures were taken along one root at described positions, and in two roots per genotype, shown are representative pictures for each position **(d)**. *Stage 0* no detectable plasma membrane detachment, extracellular vesicles or cell-wall thickening; *Stage 1* plasma membrane detached, extracellular vesicles and beginning of cell-wall thickening; *Stage 2* wild-type CS widening: secretion stops and PM is attached to modifed cell-wall in the centre; laterally, the PM is detached and extracellular vesicles are present; *Mutant stage 2 (2^m^) caspQ* foci grow: secretion continues in centre, cell-wall thickens inwards, no PM attachment detectable. *Stage 3* mature wild-type CS: no detectable secretion, PM is attached to cell wall and CS is thin and wide. *Mutant stage 3 (3^m^)* mature *caspQ* foci: secretion stops, cell wall is very thick but PM remains detached. Quantification of all stages from two roots per genotype, 8-10 cells per root **(e)**. **f-g.** Tomograms of *caspQ* foci after chemically fixation **(f)** and high-pressure freeze-substitution **(g)**. Chemical fixation induces plasmolysis, allowing visualization of PM detachment and extracellular vesicles, while in high-pressure freezing apoplastic space is compressed and vesicles are still present but squeezed. Inside the *caspQ* structure, thin transversal aligned lamellae are detected by both methods. **h.** Representative electron micrographs with KMnO_4_ lignin staining from wild-type and *caspQ* grown in mock or in presence of lignin biosynthesis inhibitor PA. Scale bars 1 µm.

We additionally noticed that this *caspQ* cell wall domain displayed a heterogenous structure after KMnO_4_ staining: lighter, unstained cell wall impregnations alternating with strongly stained, lignified regions (Figure 2f, movie 1). These lamellae-like structures were interspersed within the lignified wall and especially evident in tomograms. The tomographic reconstructions further revealed that plasma membrane detachment was continuous in z-direction and that the extracellular membranes corresponded to many small vesicles. In the cell wall, the lamellae appeared flat and arranged in a parallel fashion throughout the section.

Chemical fixation during TEM preparations always results in some degree of plasmolysis, which conveniently allows to visualize membrane attachment below the CS - and thus highlighting absence of attachment in *caspQ* (Figure 2f, movie 1). We nevertheless wanted to observe *caspQ* membrane and cell wall in a turgid cell, without chemical fixation artifacts. We therefore generated TEM tomograms after cryo-fixation (Figure 2g). Here, we could see CSD membrane was turgid and not invaginated, but still separated from the wall proper by a compacted matrix containing extracellular membranes, appearing as bigger tubules or vesicles (movie 2). Therefore, both EM methods detect the presence of EVBs and membrane separation in *caspQ.* Next, we were interested in understanding the increased heterogeneity of KMnO_4_ staining and the interspersed lamellae in *caspQ*, which suggested the presence of additional polymers. We first blocked monolignol production by treating seedlings for 24h with piperonylic acid (PA). This efficiently inhibited lignin deposition in *de novo* grown root sections, as evidenced by disappearance of KMNO_4_ staining (Figure 2h). Yet, *caspQ* still displayed abnormal cell wall thickening, to the same extent as in untreated samples. These results indicate that the mutant cell wall thickening represents the deposition of a non-lignin cell wall matrix, which then becomes impregnated with lignin. We then wanted to see whether the lamellar structures observed in the mutant are due to abnormal, “mixed” deposition of suberin and lignin. We therefore generated a pELTPxve>>GELP12 line in the *caspQ* mutant background. GELP12 is a putative suberin-degrading enzyme that has previously been shown to abrogate suberin accumulation in the endodermis (Ursache et al., 2021b) and we could observe the same effect in *caspQ* (Supplementary Figure 2.3a). Nevertheless, lamellar structures in *caspQ* were found to persist in these lines, suggesting that they are not made of suberin (Supplementary Figure 2.3b). We conclude that cell wall and membrane ultrastructure are dramatically affected in *caspQ,* with highly active and persistent secretion, associated with extensive and disorganized cell wall deposition, consisting of lignin and other, currently unknown, matrix components. Importantly, early CS development in wild-type shows a similar appearance of EVBs, but only in a restricted and transient fashion.

### caspQ foci overgrowth is due to SCHENGEN pathway activation

The SGN3 receptor pathway monitors CS integrity, by detecting diffusion of stele-derived ligands into the cortical space. Complete absence of SGN3 signaling in SCHENGEN pathway mutants leads to incomplete fusion of the CS. Chronic inability to fuse the CS in other mutants, by contrast leads to prolonged SCHENGEN pathway stimulation, causing compensatory responses, such as cell corner lignification and early suberization (Fujita et al., 2020). To test whether the observed *caspQ* phenotypes are due to SGN3-pathway activation, we targeted the SGN3 locus by CRISPR-Cas9 in the *caspQ* mutant. As expected, mutating *sgn3* removed cell corner lignification (Figure 3a) and early suberization (Supplementary Figure 3.2) of *caspQ*. Yet, in addition, *caspQ sgn3* severely reduced the number and overgrowth of the central lignin foci (Figure 3b-d). They appeared less numerous, smaller and weakly stained in *caspQ sgn3* than in *caspQ* (Figure 3a,b, Supplementary Figure 3.1a). EM ultra-structure confirmed fluorescent lignin staining, since we observed *caspQ sgn3* lignin foci less frequently and, when present, they appeared thinner and lacked much of the enhanced wall deposition seen in *caspQ* mutants (Figure 3c; Supplementary Figure 3.1b). Moreover, *caspQ sgn3* revealed an additive phenotype of smaller and less abundant central lignin foci than either *caspQ* or *sgn3* single mutants. We interpret these foci as CS-initiation sites that are positioned and formed independent of CASPs and SGN3. CASPs and SGN3 then act together to promote the growth and fusion of these sites into a final fused strip.

**Figure 3:**
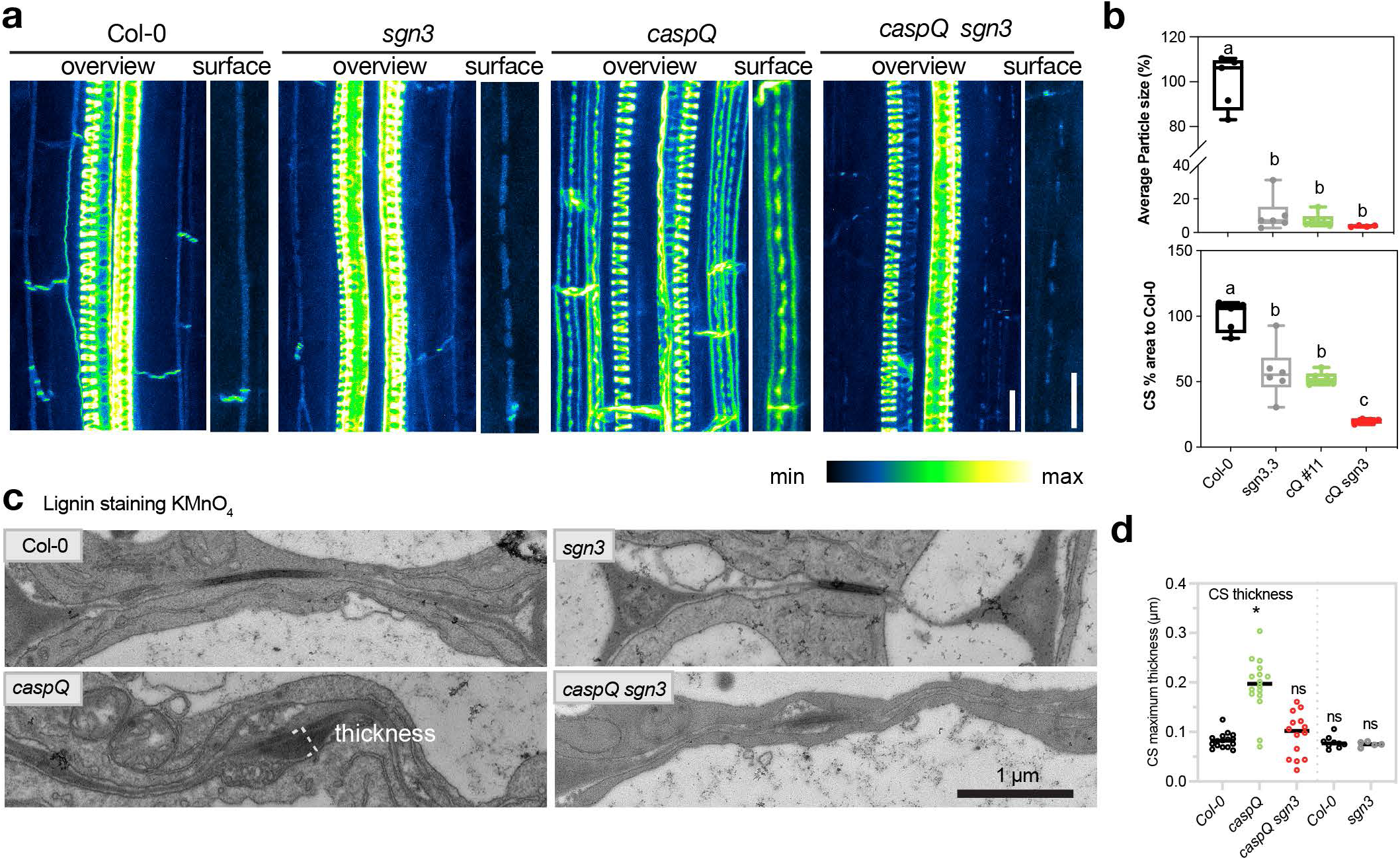
Outgrowth of caspQ lignin foci is SGN3-dependent. **a.** Maximum intensity projections of lignin stained 5-day old roots at endodermal cell number 24 in wild-type, *sgn3*, *caspQ* and *caspQ sgn3*. Scale bars 10 µm. **b.** Particle analysis of CS surface as described in Materials and Methods, with variables Average particle size (%) and CS % area to Col-0 measured per 46 µm of CS surface. Boxplots and data points (n=6) and significant differences to wild-type shown with different letter by ANOVA, Tukey’s test *p*-value < 0.05. **c.** Electron micrographs of KMnO4 lignin-stained CS of 5-day old roots at 1.5 mm from root tip in Col-0, *sgn3*, *caspQ* and *caspQ sgn3* and respective quantification of CS maximum thickness stained by KMnO4 **(d).** Shown is the mean and data points from two experimental sets (separated by dashed line, first set two roots, n∼16 CS measurements per genotype; second set one root per genotype n∼8 CS measurements). Statistical differences to wild-type by ANOVA, Dunnett’s test *p*-value < 0.05.

### The lignin machinery still accumulates in central foci in caspQ

We have previously proposed a model whereby CASPs act as scaffold proteins, bringing together a membrane-localized NADPH oxidase (RBOHF), peroxidases, such as PER64 and other lignin-polymerizing proteins for efficient and localized lignin formation (Lee et al., 2013). The full knock-out of CASP activities in the *caspQ* mutant now allowed us to directly test whether CASPs are required for localization of the known CS localized lignin-polymerizing components. We therefore introduced marker lines for the *RESPIRATORY OXIDATIVE BURST F* (*pRBOHF:RBOHF-mCherry*), *ENHANCED SUBERIN 1* (*pESB1:ESB1-mCherry*) and *PEROXIDASE 64* (*pPER64:PER64-mCitrine*) (Figure 4a-c). To our surprise, all markers accumulated in centrally aligned microdomains in caspQ. RBOHF:mCherry accumulated in a central position. However, due to its low expression and weak accumulation throughout the plasma membrane, we could not generate a surface view to resolve whether it accumulates in dot- or band-like pattern (Figure 4a). Likewise, ESB1 and PER64 secretion to the central domain was not affected in *caspQ*, but in the surface view both enzymes accumulated in dotty-pattern perfectly co-localizing to the central lignin foci of *caspQ* (Figure 4b,c). Next, we looked at ROS production using CeCl_3_ staining and EM. Upon ROS presence, CeCl_3_ produces a precipitate detectable in EM (Bestwick et al., 1997; Lee et al., 2013). In wild-type, precipitates are exclusively detected on the cortex side of the CS, presumably because CeCl_3_ cannot penetrate through the CS to form precipitate on the other side. Localized ROS production is boosted by the SGN3-pathway through SGN1 kinase-mediated activation of RBOHF (Fujita et al., 2020). Precise co-localization of all these components at the CS ensures that ROS production occurs in the defined space of the growing CS. Mutants defective in CS formation display enhanced and de-localized ROS production, due to an enhanced SGN3-pathway stimulation (Fujita 2020). *caspQ* presented CeCl_3_ precipitates throughout the whole central plasma membrane, surrounding the central foci and spanning towards cortex and pericycle sides (Figure 4d). This indicates that, while CASPs are not required for localizing ROS production to the initial central foci, they appear to be needed to restrict and confine ROS production within the foci. Nevertheless, it is difficult to distinguish whether enhanced CeCl_3_ precipitates are due to stronger ROS production or increased penetration of CeCl_3_. As for ROS production, CASP proteins are not required for correct accumulation of lignifying enzymes, but rather to restrict and organise their activity within the initial foci.

**Figure 4:**
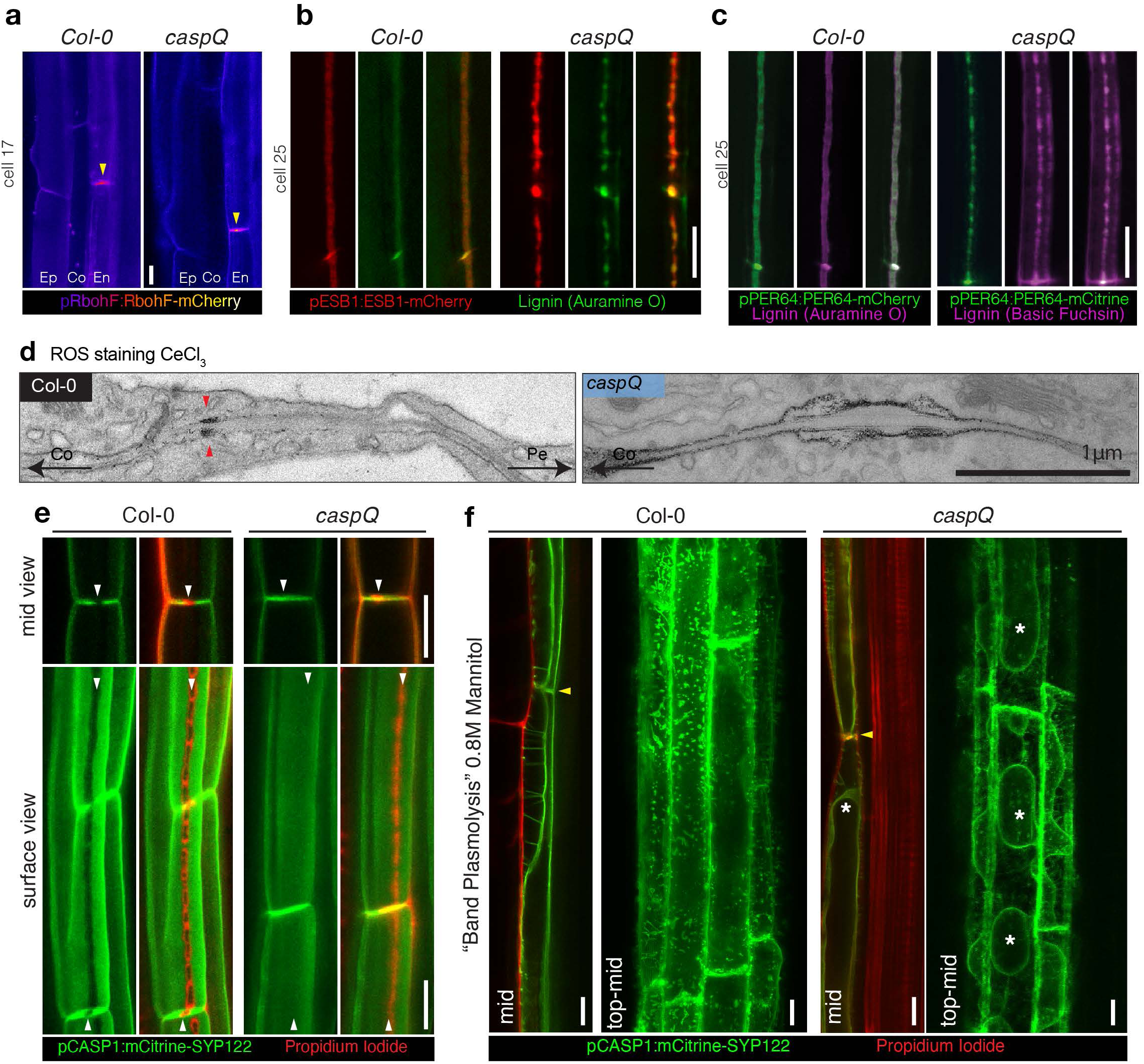
CASPs are not required for localized lignification but for CSD protein exclusion and cell-wall adhesion. **a-b.** Localization of CS-lignifying enzymes in endodermis of 5-day-old seedlings of Col-0 and *caspQ*: RBOHF-mCherry at cell 17 accumulation in CS (yellow arrow) Ep, epidermis; Co, cortex; En, endodermis **(a)**; ESB1-mCherry and Auramine O stained CS-lignin **(b)** and PRX64-Cherry with Auramine O stained CS-lignin (Col-0) and PRX64-mCitrine with Basic Fuchsin stained CS-lignin (*caspQ*). **d.** Electron micrographs from CeCl_3_ ROS-stained roots of Col-0 and *caspQ*. Note the cortex side-stained CS in Col-0 (red arrow), in contrast to the broader and plasma-membrane associated staining in *caspQ.* **e.** Plasma-membrane marker pCASP1::mCitrine-SYP122 is excluded from CSD labelled by the Propidium Iodide-stained CS in Col-0 (see white arrows). In contrast, in *caspQ* pCASP1::mCitrine-SYP122 localizes to the CSD labelled by PI (white arrows). **f.** Plasma membrane marker pCASP1::mCitrine-SYP122 and PI-stained cell-walls after plasmolysis in seedlings mounted in 0.8M Mannitol solution. For Col-0 in endodermal mid-view, plasma membranes of neighbour cells remain attached to the CS position (yellow arrow), forming the so-called band-plasmolysis; while in a maximum projection of top to mid Z-stacks membrane appears virtually all attached to the cell wall. For *caspQ*, in both mid and top-mid views the plasma membrane is detached from the CS cell wall (yellow arrow) and protoplasts can be distinguished (asterisks). Ep, epidermis, Co, cortex, En, Endodermis, Pe, pericycle. Scale bars 10 µm (except in d. scale bar 1 µm).

### CASPs are required for forming a protein exclusion zone and plasma membrane-wall adhesion

In addition to the cell wall modification itself, the CS is characterized by the formation of a membrane domain, the CSD, that excludes most plasma membrane proteins and is tightly attached to the adjacent cell wall (Alassimone et al., 2010; Reyt et al., 2021; Roppolo et al., 2011). While CASP microdomain formation perfectly co-incides with the appearance of protein exclusion and membrane attachment, it could not be established whether CASPs are indeed required for these features. To test this, we introduced the plasma membrane marker *pCASP1::mCitrine-SYP122* in *caspQ* and found that the protein exclusion zone was completely absent. In wild-type, CSD exclusion visualised by mCitrine-SYP122 can be easily monitored by complementary localization with Propidium Iodide (PI) that stains CS in its early stages. In wild-type, mCitrine-SYP122 was strictly complementary to PI-stained CS (Figure 4e). By contrast, there was clear overlap for mCitrine-SYP122 with the strongly-PI stained, central lignin foci of *caspQ* (Figure 4e). Next, we tested whether CSD attachment to the cell wall was affected in *caspQ.* When mounted in a hyperosmotic 0.8M Mannitol solution, the plasma membrane of wild-type seedlings retracts from periclinal walls but stays attached to the CS ring in the anticlinal walls, generating what is called a “band plasmolysis” (Behrisch, 1926) across different cells in a mid-view, but no obvious plasmolysis in a top-to-mid view due to the lateral anchoring to the CS (Figure 4f). In contrast, in *caspQ* we observed a plasmolysis typical for other cell types, with membrane detachment and protoplasts appearing in all views similar to one displayed in *myb36,* which lacks the entire CS differentiation program (Figure 4f; Supplementary Figure 4.1). These finding are consistent with our EM observations, where *caspQ* displayed strong secretion, associated with plasma membrane detachment or invagination, below the lignified cell wall in early stages. In later stages, association to the cell wall resembles that of a normal plasma membrane, with weak, fixation-induced detachments (Figure 2a-d). Altogether, these results show CASPs are required to form the two defining features of the CSD, i.e. protein exclusion and membrane-cell wall attachment.

### A minimum of three CASPs are required of complementation of the quintuple mutant

Having obtained a “blank slate” in the endodermis with respect to CASP activity also finally allowed us to test whether individual CASPs have distinct or redundant function. Analysis of the series of single, double, triple and now quintuple mutants (Figure 1) already suggested partial redundancy between CASPs. We decided to use a “bottom-up” approach, i.e. to reconstitute CASP function in *caspQ* by introducing CASPs one by one. By observing localization of GFP fusions of CASP1-5 in wild-type with greater detail than previously, we observed that each CASP displayed a slightly different localization with respect to the Basic Fuchsin-stained CS. CASP1 appeared broader and with preferential localization at the edges of the strip (Reyt et al., 2020); CASP2 and CASP5, by contrast, appeared in more central, dotted domains; whereas CASP3 and CASP4 were localized in a more homogenous fashion (Figure 5a). This reinforced the idea that CASPs cannot be fully redundant to each other. Indeed, when introduced into *caspQ*, none of the single CASPs was able to restore the mutant phenotype (Figure 5b,c). Lignin in all lines displayed the typical ectopic cell corner lignification, indicative of a defective CS. However, the central lignin foci appear different in different CASP complementing lines. In CASP1-, CASP3- and CASP4-GFP *caspQ* complementing lines, we could see an improvement of the lignin foci of *caspQ*, insofar as there was a more fused lignin distribution, with less individual foci; CASP2 and CASP5, by contrast failed to improve the *caspQ* lignin foci (Figure 5c,d). Inspection of CASP-GFP accumulation along the root revealed that the proteins were generally unstable and degraded as singles and showed a more heterogenous sub-cellular localization with more vesicles and a more delocalized plasma membrane pattern. CASP5-GFP showed an intriguing particularity in that it appeared stuck in the ER and was never observed at the central plasma membrane domain in *caspQ* mutants (Figure 5c,f). However, a CASP5-mCherry could be “mobilised” by co-expression with CASP1-GFP, in which case it reached the central plasma membrane domain, where it co-localized with CASP1-GFP (Figure 5f).

**Figure 5:**
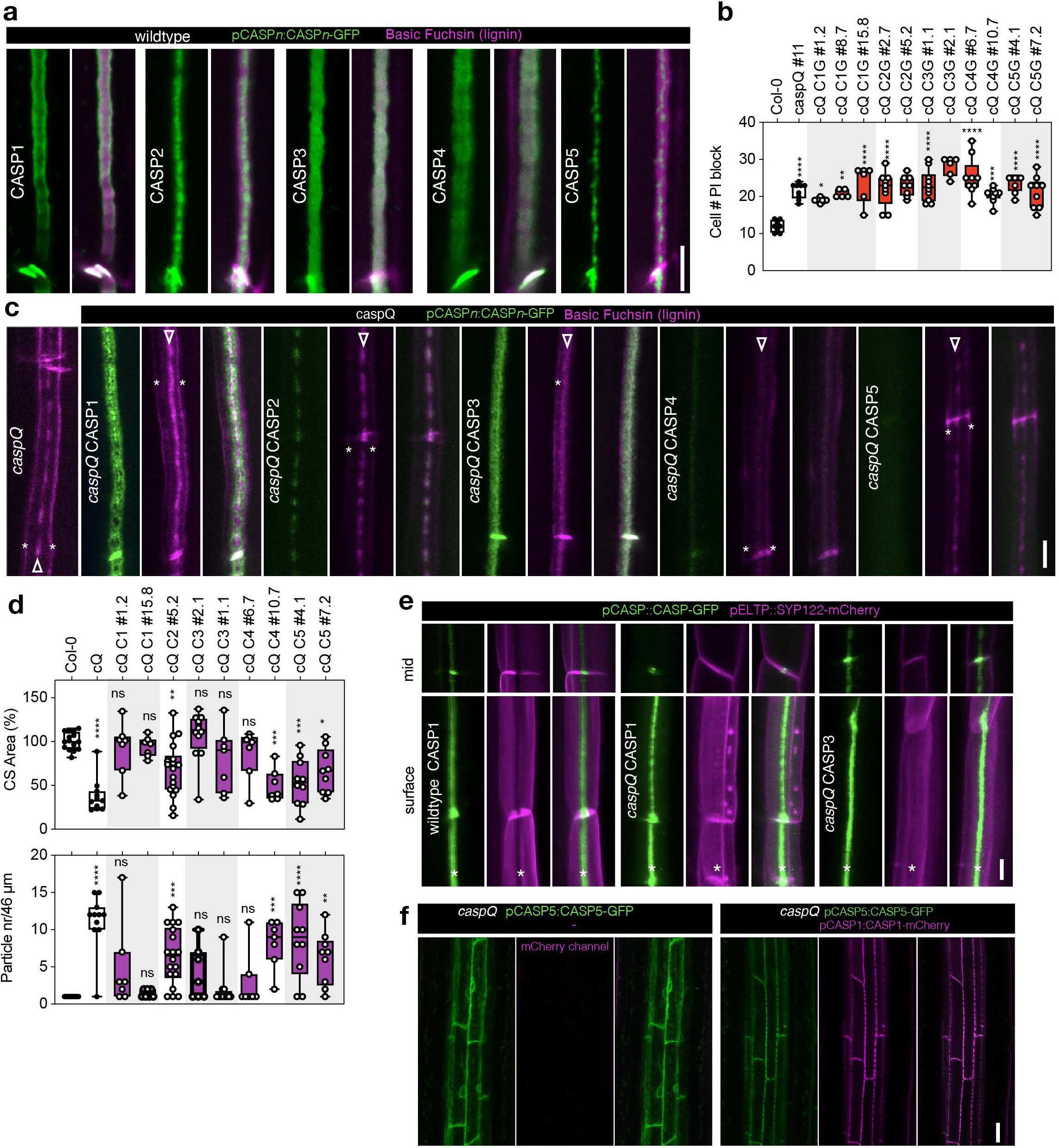
CASP1-CASP5 diverged in nanodomain localization and function to cooperate in Casparian Strip growth. **a.** CASP1- to CASP5-GFP (green) nanodomain localization in CS surfaces stained with Basic Fuchsin (magenta) at endodermal cell number 15 of wild-type seedlings (pCASPn:CASP-GFP lines). **b.** Propidium iodide uptake assay in Col-0, *caspQ* and single *CASP1-5* complementation *caspQ* + pCASPn:CASP-GFP, n=6-8 individuals, from at least two independent T3 homozygous lines per gene. Statistical differences by ANOVA, Dunnet’s test comparisons to Col-0 (p<0.05). **b.** Detailed surface view of lignin staining with Basic Fuchsin and CASP1-5:GFP localization in *caspQ* and *caspQ* single *CASP1-5* complementation lines at endodermal cell number 25. Lignin stain at cell corners typical of CS-defective mutants (asterisks), and at presumptive CS (arrowheads). **d.** Particle analysis in CS surface pictures as in (c) for Col-0, *caspQ* and complementation *caspQ* + pCASPn:CASP-GFP, n=8-12. Analysis done as in Fig. 1, yield CS Area (%) relative to Col-0 (upper graph), and particle number along ∼46 µm of CS length. Statistical differences by ANOVA, Dunnett’s test comparisons to Col-0 (p<0.05). **e.** CSD exclusion zone (asterisk) in wild-type and *CASP1-5* complementation *caspQ* + pCASPn:CASP-GFP visualized with plasma-membrane marker pCASP1:SYP122-mCherry. Neither CASP1-GFP, nor CASP3-GFP presence is sufficient to create a CSD exclusion for SYP122-mCherry in *caspQ* background. **f.** pCASP5:CASP5-GFP in *caspQ* is retained in ER, but is able to reach CSD when co-expressed with pCASP1:CASP1-GFP in *caspQ*. Scale bars 5 µm (a, c, e) and 20 µm (f).

By crossing CASP1/3-GFP *caspQ* to a *caspQ* containing a plasma membrane marker (pELTP::mCherry-SYP122), we found that neither CASP appeared to be able to form a protein exclusion zone, despite CASP1/3 appearing to be correctly targeted to the central plasma membrane domain, suggesting that interaction between several CASPs is needed to generate a CSD exclusion zone (Figure 5e). By crossing *caspQ* CASP1-mCherry with *caspQ* CASP2 to 5-GFP we could demonstrate that also CASP double combination could not effectively rescue the *caspQ* PI phenotype (Supplemental Fig 5.1a). In order to identify a minimal CASP combination for rescue of *caspQ,* we generated a combinatorial Gateway collection of CASP genomic entry clones for triple Gateway. In this system, we included whole genomic regions to ensure proper transcription start and termination, generating single, double and triple combinations. We then screened for PI block of individual T1 seedlings for each construct. We confirmed that single and double combinations cannot rescue *caspQ* (Supplementary Figure 5.1b). Yet, we found that one triple combination (CASP1, CASP3 and CASP5) could efficiently complement *caspQ* in most independent T1 individuals. In other combinations, we could also detect complementation with some individual lines (Supplementary Figure 5.1b). However, these combinations varied in complementation efficiency depending on the CASP position in the construct (e.g. CASP1+CASP3+CASP2, better than CASP1+CASP2+CASP3), indicating positional effects within our multi-gene constructs. We confirmed this T1 complementation analysis in a selected number of T2 lines (Supplementary Figure 5.1c). Despite these positional effects, our multiple efforts to find a minimal CASP function suggest that a minimum of three CASPs is necessary to restore endodermal *caspQ* function, with a combination of CASP1, CASP3 and CASP5 being the most efficient in our hands.

In summary, the five endodermal expressed CASPs of *Arabidopsis* present an evident degree of diversification, both in their nanodomain localization within the CS, as in their ability to partially rescue *caspQ* lignin deposition. Yet, the combined activity of at least three CASPs is required to restore a fully functional strip in *caspQ*.

### CASPs evict EXO70A1 from CS initiation sites

Immediately preceding the appearance of CASPs, EXO70A1 targets the exocyst complex transiently to the position of the incipient CSD, its knock-out leading to complete delocalisation of CASP secretion (Kalmbach et al., 2017). Since *caspQ* appears to lack a CSD, but is nonetheless able to form correctly localized, albeit aberrantly structured, lignin foci, we investigated how the exocyst complex localized in the mutant. We monitored the appearance of EXO70A1, the subunit responsible for exocyst localization and assembly (Kalmbach et al., 2017; Synek et al., 2021), in wild-type and *caspQ* using an EXO70A1 line with endodermis-specific expression (*pSCR::EXO70A1-mVenus*). Endodermis-specific expression of EXO70A1 was used to improve imaging and allow surface observations. As reported for the native promoter line, pSCR::EXO70A1 accumulated transiently at CSD in wild-type (Kalmbach et al., 2017) (Figure 6a,b). Despite being weakly expressed from the early-endodermis *SCR* promoter, we were able to precisely image EXO70A1 dynamic at the endodermal surface. Interestingly, when introduced in a *pCASP1::CASP1-mCherry* background, a mutually exclusive localization of EXO70A1-mVenus and CASP1:mCherry in wild-type was observed (Figure 6c). EXO70A1 accumulated in the gaps between CASP1-mCherry at the string-of-pearls stage and later surrounded the strip, as if EXO70A1 was excluded from where CASP domains have formed, accumulating instead in spots where CASP1 is still to be delivered, i.e. the gaps of string-of-pearls and the edges during CS widening. This dynamic was consistent with the mid-section views, where EXO70A1 precedes CASP1 appearance, but then disappears from CASP1-mCherry containing spots, such that the two proteins hardly co-localize. These observations suggested that CASP localisation and EXO70A1 localisation are somehow mutually exclusive. Intriguingly, in *caspQ*, the same initial localization of EXO70A1 was observed. Yet, EXO70A1 then showed a systematically increased persistence at the central foci in the mutant (Figure 6a,b). Moreover, when we observed EXO70A1 in surface views together with PI as lignin stain, EXO70A1 was again perfectly complementary to the PI-stained domains at the string-of-pearls stage in wild-type. Again, this pattern was completely reversed in *caspQ,* where EXO70A1 localisation strongly coincided with the PI-stained lignin foci (Figure 6d). We next introduced pESB1:ESB1-mCherry to monitor a *bona fide* CS marker that could be used in the two backgrounds. Here again, we observed a negative correlation of ESB1 and EXO70A1 in wild-type and a strictly positive correlation of the two proteins in *caspQ* (Figure 6e). We were then interested in localization of EXO70A1 in *sgn3*, a mutant also with interrupted CS but with improved CS coverage than *caspQ* (Figure 1c). When introducing *pSCR::EXO70A1-mVenus* in *sgn3* pESB1:ESB1-mCherry lines, we could see an intermediate phenotype to wild-type and *caspQ*, i.e. EXO70A1 was partially displaced from CS islands to their edges, but not localizing strictly to the gaps as in wild-type (Supplementary Figure 6.1). This suggests that SGN3 contributes to the dynamic EXO70A1 relocation to new domains, which could be one of the reasons for the slow growth and incomplete fusion of CASP microdomains in *sgn3*.

**Figure 6:**
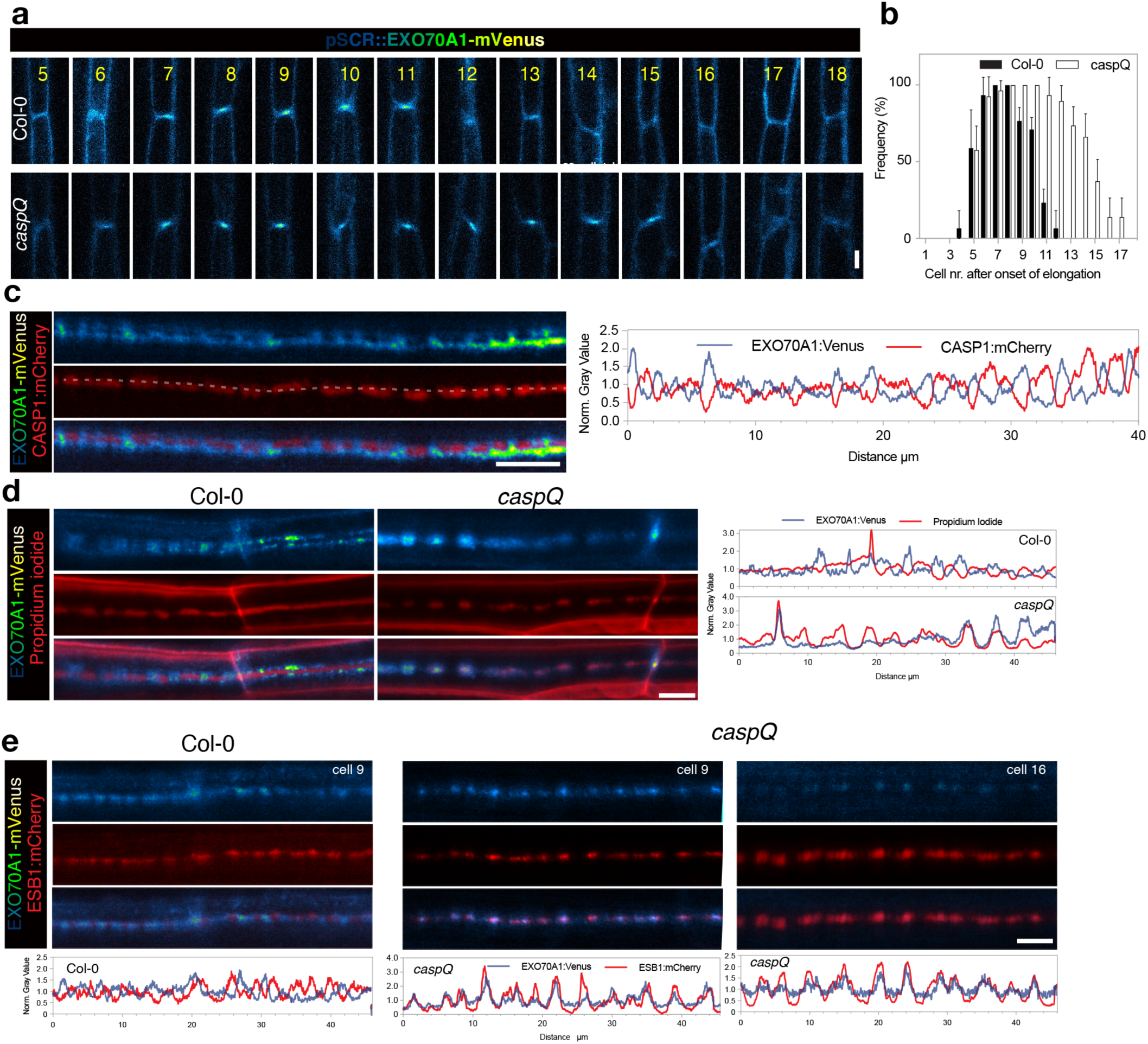
CASPs are necessary for EXO70A1 removal and progression in growing Casparian Strip. **a-b.** Dynamic accumulation of EXO70A1-mVenus at CSD, inspected along consecutive endodermal cells (5 to 18) in Col-0 and *caspQ*, confocal picture from representative individuals **(a)** and quantification in n=8-10 individuals **(b)**. **c.** Mutual exclusion of EXO70A1-mVenus and CASP1:mCherry in CS surfaces from wild-type seedlings at cell number 8. Shown is the profile for each channel obtained from same picture, drawing a line of 20pt thickness (exemplified by dashed gray line). **d.** Mutual exclusion (wild-type) and co-localization (*caspQ*) of EXO70A1-mVenus and Propidium iodide in CS surfaces in cells number 8-9 with respective profiles, obtained as in c. **d.** Mutual exclusion (wild-type, cells number 8-9) and co-localization (*caspQ,* cells number 9 and 16) of EXO70A1-mVenus and ESB1-mCherry in CS surfaces with respective profiles, obtained as in c. Scale bars 5 µm.

In summary, the mutually exclusive localization of CASP and EXO70A1, together with the fact that EXO70A1 persists at the lignin foci of *caspQ,* indicates that CASPs act to evict EXO70A1 from the forming CS domains, thereby forcing its re-localisation to the edges and into the remaining gaps.

### CASP1-proximity labelling identifies novel, CS-associated Rab GTPases

CASPs are very small transmembrane domain proteins with limited surfaces for biochemical functions. In our efforts to understand CASP function, we envisioned that identifying CASP-interactors could help us understanding how these small proteins play such a critical role in CS development. In previous attempts to identify interactors by co-immunoprecipitation using CASP1-GFP as bait, we could only confidently identify CASP3 (Roppolo et al., 2011). We attributed these difficulties to CASP1-GFP low solubility from low-speed pellets, due to the strong attachment of the CSD to the CS cell-wall. To tackle this problem, we decided to employ proximity labelling fusing the biotin ligase, TurboID, to CASP1 (pCASP1::CASP1-sGFP-TurboID, short CASP1-tID). Proximity labelling employs degenerated biotin ligases that convert applied biotin into highly reactive biotin-AMP that diffuses out of enzyme core and covalently attaches to proximal proteins within a defined radius (Branon et al., 2018; Mair et al., 2019). The advantages of this method for our purpose were the broader detection of both strong and weak interactors, and, as biotinylation occurs *in vivo*, the possibility to do stringent washes to reduce contaminants and increase solubility of tightly CS-associated proteins. Since the CS is not a membrane confined compartment, we had to filter for abundant cytosolic or plasma membrane proteins that would be labelled just by “passing-by”. We therefore generated two TurboID controls expressed under the same *pCASP1* promoter: a cytosolic TurboID (*pCASP1::sGFP-TurboID*, cyto-tID) and a plasma membrane (*pCASP1::sGFP-TurboID-SYP122*, PM-tID). Thanks to the GFP moiety we could ascertain the expected endodermal expression, as well as the correct localization of the fusion proteins to cytosol, plasma membrane and CS domains (Figure 7a) and protein integrity by Western Blot (Figure 7b). After biotin feeding for 24h, the three lines showed a detectable pool of biotinylated proteins of diverse sizes and abundances, with the strongest band corresponding in size to predicted self-biotinylation of the GFP-TurboID fusion proteins (Figure 7b). We then purified these proteins and conducted LC-MS/MS-Spec to identify the biotinylated proteins in each TurboID samples with pCASP1::CASP1-GFP plants as negative control and three biological replicates per sample, following established protocols (Branon et al., 2018; Mair et al., 2019). We identified ∼3800 proteins in each TurboID sample and 1800 in the negative control. In clustering and PCA analyses, the biological replicates cluster together, and CASP1-tID sample was closer to the PM-tID than the cytosolic control (Figure 7c). Protein abundances were calculated using iBAQ, ‘intensity-based absolute quantification’, i.e. protein intensities (the sum of all identified peptide intensities) divided by the number of theoretically observable peptides (calculated by *in silico* protein digestion), and values were log2-transformed for visualization purposes (Schwanhäusser et al., 2011). As expected, self-biotinylated GFP-tID and respective fusion proteins scored among the most abundant proteins in TurboID samples. Moreover, GFP-tID abundance was of similar intensity (log2iBAQ from 8 to 10) among replicates and tID samples, implying labelling and purification yields were comparable, allowing to confidently compare iBAQ values between the distinct transgenic lines.

**Figure 7:**
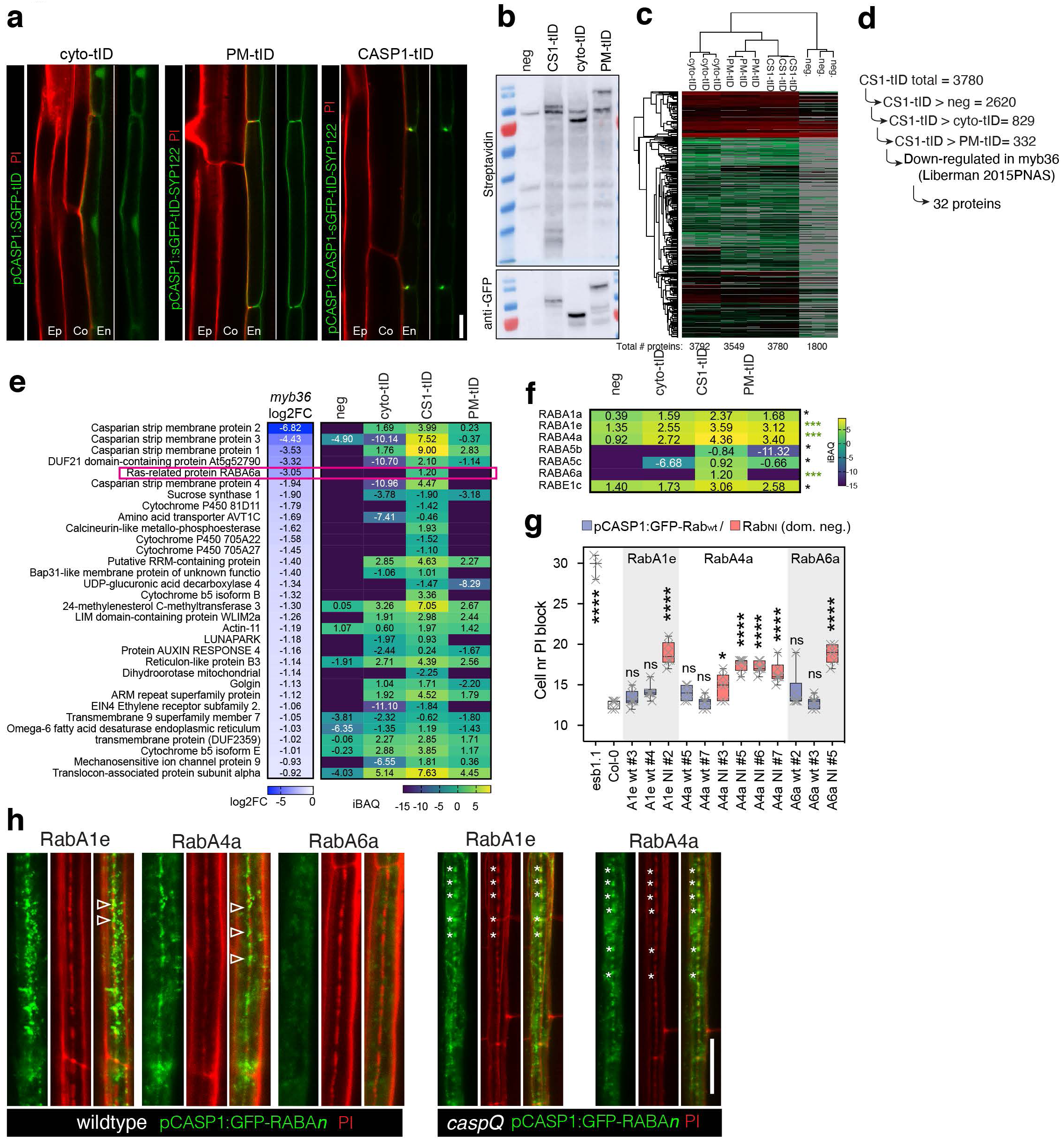
CASP1-proximity labelling identifies RabAs with CASP-dependent CS-localization and function in CS integrity. **a.** Endodermal expression and sub-cellular localization of GFP-turboID protein fusions: cytosolic (*cyto-tID*, pCASP1:sGFP-turboID), plasma membrane (*PM-tID*, pCASP1:sGFP-turboID-SYP122) and CS-localized (*CASP1-tID*, pCASP1:CASP1-sGFP-turboID). **b.** Western Blots of roots of *cyto-tID*, *PM-tID* and *CASP1-tID*, negative control (pCASP1:CASP1-GFP) grown for 6d in normal media and additional 1 day in biotin (50µM) containing plates; showing biotinylation pattern and presence of intact turboID fusion constructs. Samples refer to input before purification. **c.** Cluster Analysis of LC/MS-MS identified proteins from streptadividin purified root extracts as in (b) done in Perseus, from three biological replicates of *cyto-tID*, *PM-tID*, *CASP1-tID* and negative control. **d.** Sequential pairwise analysis (t-Test, Bonferroni adjusted *q*-value <0.05) reveal 332 significantly enriched proteins in *CASP1-tID* relative to all other samples. From those 332 proteins, 32 are significantly down-regulated in mutant of CS-transcriptional regulator *myb36.* **e.** Transcriptional downregulation in *myb36* (log2 Fold Change, Liberman 2015 PNAS) and protein abundance in turboID experiment (as in b and c, iBAQ values) of the 32 proteins significantly enriched in CS1-tID samples and significantly downregulated in *myb36,* highlighting Ras-related small GTP-ase, RabA6a. **f.** Significantly enriched Ras-related small GTP-ases in *CS1-tID* sample, *** p<0.01 and * p<0.05. **g.** Propidium uptake assay of endodermal specifically expression of the three most enriched RabAs (A1e, A4a and A6a, enriched in CS1-tID p<0.01) as wildtype sequence and as dominant negative N-I mutants, Col-0 and *esb1-1* as reference. Statistical differences by ANOVA, Dunnett’s test comparisons to Col-0 (p<0.05). **h.** Localization of the three GFP-RabAs in wiltype and *caspQ* (with exception of RabA6a, where no lines with detectable signal could be retrieved) in cell number 9 stained with Propidium Iodide. Not the presence of GFP-Rab signal intercalated with PI-stained CS string-of-pearls (arrowhead) in wiltype, in contrast with perfect co-localization of GFP-Rab with PI stained central foci of *caspQ* (asterisks). Representative profiles of GFP-RabA4a and PI in Col-0 and *caspQ* CS-surfaces. Scale bars 10 µm.

**Figure 8:**
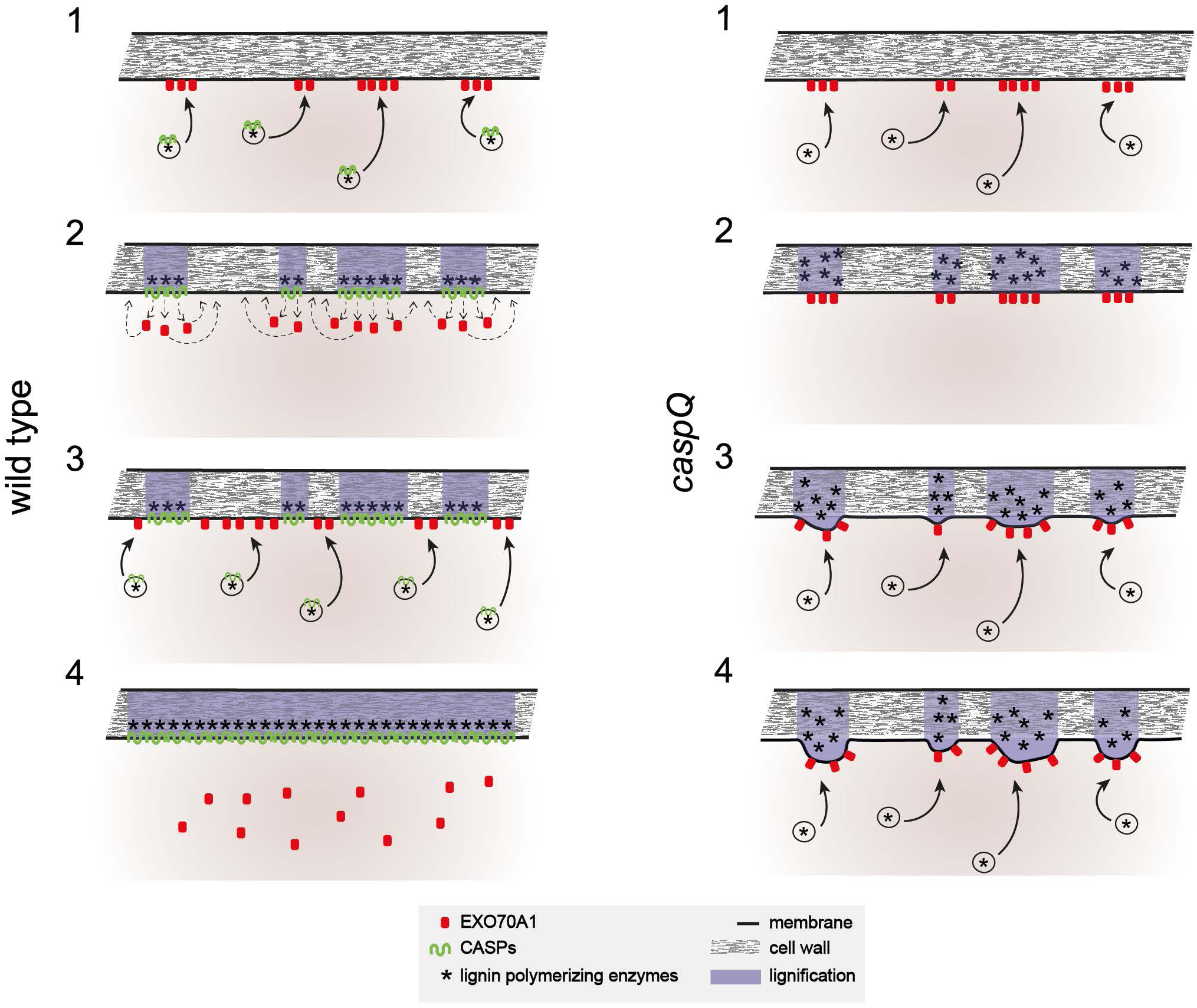
Model of CASP action in the endodermis. *In presence of CASPs (**wild-type**)*: Vesicles carrying CASPs and lignin polymerizing enzymes are specifically targeted to EXO70A1 landmarks at the endodermal plasma membrane (1). Targeting specificity is possibly mediated by the presence of specific RabAs on the vesicles (not shown). CASPs form stable microdomains by polymerization, organizing lignin polymerizing enzymes in the cell wall and evicting EXO70A1 on the cytosolic membrane face. EXO70A1 re-locate to parts of the central region that are still free of CASPs (2). New CASP carrying vesicles deliver CASPs to the remaining gaps between microdomains marked by EXO70A1 (3), eventually fusing individual microdomains into a contiguous band and thus sealing the cell wall space (4). *In absence of CASPs (**caspQ**)*: Vesicle carrying lignin polymerizing enzymes are correctly targeted to EXO70A1 landmarks (1), forming initially normal lignin foci in the correct position (2), absence of CASPs allows EXO70A1 landmarks to persist in the same initial position, further attracting vesicles and lignin polymerizing activity to the same foci, leading to their thickening (3). Continued, persistent secretion to the same foci eventually leads to strongly thickened, aberrantly structured lignin foci that are unable to fuse with each other and do not seal the cell wall space (4).

We then checked if known players involved in CS formation were detected and how they behave in different turboID samples (Supplementary Figure 7.1). Besides CASP1, we detected CASP2, CASP3 and CASP4, specifically enriched in CASP1-tID. CASP5, the smallest CASP protein, although exclusively detected in CASP1-tID, did not pass the quality criteria for LC-MS/MS-identification, i.e. to be identified by more than one peptide, thus could not be included in the analysis. Interestingly, CASPL1A1 was significantly enriched in CASP1-tID and PM-tID samples, supporting previous report of CASPL1A1 localisation to the CSD when expressed under CASP1 promoter (Roppolo 2014), but also a broader PM-localization in native conditions, fitting to our current genetic analysis where CASPL1A1 knockout does not contribute to CS formation.

Members of the EXOCYST complex were also detected in turboID samples, but with no preference for CASP1-tID sample, in accordance with the reported transient accumulation to the CSD and otherwise broad expression and localization to the plasma membrane (Kalmbach et al., 2017; Synek et al., 2021). Notably, we identified one or two members of each subunit of the complex (SEC3,5,6,8 and 15,84,70), with exception of SEC10 for which no isoform was identified. All members were more abundant in PM-tID and variably abundant in cyto-tID vs. CASP1-tID, e.g. EXO70A1 iBAQ scored highest in PM-tID and then in CASP1-tID; SEC6 iBAQ scored highest in PM-tID and then in cyto-tID. These observations fit the complex’s broad function in membrane trafficking and its dynamic cytosolic and membrane-associated localization (Ahmed et al., 2018).

From the SCHENGEN signalling pathway, we could not identify SGN3, but did identify SGN1, RBOHF and RBOHD. Importantly, the three membrane proteins are absent from cyto-tID, but appeared with similar abundance in CASP1-tID and PM-tID samples. This is expected for SGN1 and RBOHD neither of which localises specifically at the CSD. RBOHF by contrast shows a clear accumulation at the CSD, yet with detectable lower levels everywhere in the plasma membrane (Figure 4a). Thus, the absence of increased peptide counts in the CASP1-tID fraction could simply be explained by the many times higher surface that PM-tID protein occupies, compared to CSD localized CASP1-tID (Fujita et al., 2020; Lee et al., 2013). Finally, we surveyed for the GO-term lignin, and could detect three CASP1-tID specific proteins, i.e. trans-cynnamate 4-monoxygenase (or C4H, Cytochrome P450 73), cytochrome P450 98A3 and Glucuronoxylan 4-O-methyltransferase 3 and/or 2, which are predicted to localize to ER and/or Golgi membranes, according to SUBA4. However, we did not detect CS-localized lignification enzymes, e.g. ESB1, PER64, UCC1 (Hosmani et al., 2013; Lee et al., 2013; Reyt et al., 2020). Indeed, their apoplastic localization separates them by the plasma membrane from the biotinylation reaction occurring on the cytosolic side of CS.

Next, we filtered for CASP1-turboID enriched proteins using sequential pairwise t-Tests. Using presence of CASP2-4 as reference, we found the most adequate cut-off for p-value at 0.05; and to be as inclusive as possible, we considered log2iBAQ differences above 0. Starting with the comparison CASP1-tID versus the negative control, from the total of 3780 detected proteins identified in CASP1-tID, we removed 1160 proteins that could either be endogenously biotinylated proteins or contaminants from the streptavidin purification. Next, from the remaining 2620 proteins, 829 were significantly enriched compared to cytosolic-tID; and finally, 332 proteins were significantly enriched compared to PM-tID (Figure 7d, Table 2).

After this filtering, CASP1 appears as the most abundant protein, followed, interestingly, by ubiquitin; CASP3 appears as the 5^th^, CASP4 the 38^th^ and CASP2 the 49^th^. In a quick survey of the list, we find several proteins associated with protein degradation. Besides Ubiquitin itself, 26S proteasome subunits, proposed and *bona fide* members of ESCRT complex, TOM-like proteins, and FREE1/FYVE (Gao et al., 2017). Another interesting category is sterol methyl esterification, with the three Arabidopsis SMTs, ranking among the ten most highly abundant proteins. Finally, many enzymes for monolignol biosynthesis or activation were found, e.g. trans-cinnamate 4-monooxygenase; and three isoforms of cytochrome B5 and one NADPH-cytochrome B5 reductase; four cytochrome P450, and three NADPH-cytochrome P450 reductases, and MSBP2 (membrane steroid-binding protein 2) a proposed scaffold for the ER-localized monolignol biosynthetic P450 enzyme complex (Gou et al., 2018)

Overall, sub-cellular localization analysis revealed a striking prevalence of ER-localized proteins in the CASP1-turboID fraction (Supplementary Figure 7.2). Another category of particular interest was “vesicle transport”. When filtering for these key words in GOBP, we found an intriguing overrepresentation of the A subclass of Rab proteins. Of the 57 *Arabidopsis* Rab-GTPases (Rutherford and Moore, 2002), we found 36 members in the whole experiment. Seven of those were specific to CASP1-tID sample (Supplementary Figure 7.3). Moreover, when additionally filtering for proteins showing decreased expression in *myb36*, we found one RabA, RabA6a, which was exclusively detected in the CASP1-tID fraction (Figure 7e,f). Moreover, five other RabAs, that are not MYB36 regulated, also showed significant enrichment in the CASP1-tID fraction (Figure 7f, Supplementary Figure 7.3). Rabs are of particular interest, because they are central regulators of vesicle fusion and membrane organisation and the RabA subclass, in particular (mammalian Rab11/25 subclass homologs) are known to be involved in late steps of secretion or endosomal recycling (Rutherford and Moore, 2002). Since activated Rab proteins are moreover known activators of the exocyst complex, we tested whether Rabs are important in CS formation by generating cell-type specific overexpressors of dominant-negative RAB variants. Indeed, dominant-negative lines of the three most significantly enriched (p<0.01) RabA homologs (A1e, A4a and A6a) showed a weak, but consistent delay in the formation of the apoplastic barrier, suggesting a role in CS formation and possibly CASP trafficking (Figure 7g). Moreover, localisation of two RabAs expressed under an endodermis-specific promoter, showed RabA-positive vesicular structures to be in close vicinity, but distinct from, initiating CS patches in wild-type. The *myb36*-regulated RabA6a signal was very weak, and we could not detect specific CS accumulation. By contrast, these RabA-positive compartments strongly overlap with cell wall foci in *caspQ* (Figure 7h). Although, based on vesicular structures and not membrane domains, this situation closely resembles that of EXO70A1 described earlier and is consistent with RabAs acting as activators of exocyst during targeting of CASP-bearing vesicle to the central domain.

## Discussion

### MARVEL protein function in eukaryotes

CASPs and CASP-LIKES represent a plant-specific branch of the MARVEL superfamily, a highly sequence divergent family of four trans-membrane spanning proteins (Roppolo et al., 2014; Sánchez-Pulido et al., 2002). Their presence not only in ophistokonts (animals and fungi), but also in plants, as well as heterokonta (oomycetes, brown algae) suggests that they are a truly ancient invention of eukaryotic cells (Roppolo et al., 2014). Interestingly, their conservation is largely restricted to their overall topology (four transmembrane domains with termini inside) and few specific features of the transmembrane domains. MARVEL proteins display low conservation and independent evolution of complex sub-families in animals and plants. *A. thaliana*, for example, has 5 subfamilies with a total of 39 members, whereas *H. sapiens* has 4 subfamilies for a total of 27 members (Roppolo et al., 2014; Rubio-Ramos et al., 2021). Nevertheless, there are tantalizing parallels in supposed functions of these proteins in different cell types. In animals, the four subfamilies are involved in seemingly divergent processes, such as regulation of tight-junction structure and functionality, regulation of pre-synaptic vesicle release, subdomain dynamics and signaling in immune cells, or cancer development (Antón et al., 2008; Raja et al., 2019; Saito et al., 2021; Wu et al., 2020). In attempts to formulate an overarching, basic cellular function, it was pointed out that animal MARVEL proteins are often found to be highly abundant, conserved (i.e. potential orthologs are found in insects and nematodes) and associated with the formation of plasma membrane subdomains and/or late secretory events. Surprisingly, in animals, even higher-order mutant and sometimes presumably full knock-outs of their respective functions, often leads to comparatively subtle phenotypes. Double mutant of occludin and tricellulin, for example, lead to changes of branching patterns of tight-junction strands, without interfering with their presence or overall functionality (Saito et al., 2021). The same applies for presumably full knock-out of synaptic MARVELs, which does not fundamentally interfere with synaptic transmission, but merely alters probability of synaptic vesicle fusion under certain conditions (Raja et al., 2019). A strong knock-down of MAL function, by contrast, leads to rather strong defects in the formation of a plasma membrane microdomain, called the immunological synapse, as well as signalling in T-cells (Antón et al., 2008). In budding yeast, a similar function in assembly of microdomains called eisosomes was observed for the MARVEL protein Nce102 (Loibl et al., 2010).

### CASP function in Casparian strip formation

Our full knock-out of the endodermis-expressed CASPs, now provides one of the clearest MARVEL-phenotype in multi-cellular eukaryotes and could serve as a paradigmatic example for MARVEL function beyond plants. Our data indicates that CASP-driven microdomain formation at the plasma membrane suppresses further secretion at a given site by excluding other PM proteins and evicting EXO70A1, the factor that is central for targeted secretion of CASP-bearing vesicles in the first place. The consequence of this negative feedback loop is to enforce continual displacement of EXO70A1 secretory foci along the median zone, leading to rapid filling of gaps in the initial string of microdomains and their eventual fusion into an uninterrupted ring. Protein exclusion and inhibition of exocytosis, and possibly endocytosis, could be a critical feature to maintain CSD identity and stability of its membrane-wall attachment, i.e. the plasma membrane of the CSD would be a relatively inert surface for trafficking and protein diffusion.

The second aspect of CASP function lies in the feature of the microdomain itself. In the absence of CASPs, two hallmarks of the Casparian strip domain, membrane-attachment and membrane exclusion zone formation are clearly absent. Thus, in plants, MARVEL proteins are not only required for functions mediated by claudins in animals, but also by adherens junction-forming cadherins. Finally, CASP domains are clearly necessary to form the homogenous, dense and spatially confined lignification that gave rise to the term “strip”, from the German “Streifen”, describing its flat appearance and well-defined width. In the quintuple CASP mutant, centrally focused delivery of the cell wall modifying enzymes is still occurring, but the resulting cell wall is clearly disorganized, of variable shape, excessive thickness and displaying currently unidentified, lamellae-like inhomogeneities within the lignified wall. Thus, although not required for enzyme localization at the cellular scale, CASPs do appear to be necessary to organize enzymatic activities at the nanoscale.

The ability to generate stable protein platforms in the PM, which can then organize extracellular activities, engage in cell wall/matrix interactions or establish cell-to-cell contacts, is critical for cells within complex cellular communities. Possibly, this ability is the common denominator of MARVEL protein function, that has been maintained and neo-functionalised in many different branches of life and is now used to support the function of a multitude of different tissues and cell types in extant eukaryotes.

### Extracellular vesicles are associated with Casparian strip formation

Finally, the strong and sustained secretion to a few median foci in the CASP quintuple mutants also revealed the presence of extracellular vesicles below the lignifying wall. Indeed, their presence in the mutants prompted us to look for EVs during Casparian strip formation in wild-type, where we found EVs to also be present, but only very transiently. The presence of EVs is intriguing, considering that current models for the apoplastic delivery of lignifying components postulate passive diffusion or transporter-mediated transport of monomers (Alejandro et al., 2012; Perkins et al., 2022) and lignifying enzymes to be transported by the canonical secretory pathway. Moreover, a recent report described very similar structures to be associated with suberisation in the endodermis (De Bellis et al., 2022). Since we have no indication that either *caspQ* lignin foci or wild-type Casparian strips contain suberin, we conclude that EVs in the endodermis are not associated only with suberisation, but also lignification and might be regarded generally as being associated with strong, focused secretion in non-elongating cells.

### RabAs as potential activators of specific exocyst complexes

The plant exocyst complex clearly appears to have evolved a divergent and more complex role in plants than the one described in animals and fungi. The presence of 23 different EXO70 subunits suggested that the plant exocyst is not a general component required for tethering of any plasma membrane bound vesicles, but that numerous, distinct exocyst holocomplexes might exist whose role could be to mediate tethering of specific vesicle subpopulation with distinct cargos, to different plasma membrane subdomains (Žárský et al., 2009). Yet, this model requires a matching complexity on the vesicle side that would allow to “read” different EXO70 landmarks, such that a given vesicle could be preferentially tethered to a specific EXO70 landmark. Intriguingly, plants present an additional significant radiation not observed in animals, the presence of 26 RabA GTPases, compared to only 3 Rab11/25 (RabA homologs) family members in mammals (Rutherford and Moore, 2002). Here, we found that CASPs are in specific, close proximity to RabA family members, that RabAs localize to the central domain defined by EXO70A1, immediately suggesting that RabA-bearing vesicles might have specific affinity for EXO70A1 marked domains. Indeed, cell-type specific, dominant-interference lines of different RabA homologs collectively showed that RabA have some function in CS formation. In animals and yeast clear evidence points to exocyst being a “Rab effector”, i.e. the exocyst being activated by Rabs (e.g. Sec4 in yeast, Rab11 in animals) (Guo et al., 1999; Zhang et al., 2004). It is therefore intriguing to speculate that plant cells could produce different classes of late secretory vesicles, discriminated by different RabA subclasses and that these vesicles have preferences for specific EXO70 subunits thus allowing for tethering of different vesicle populations to distinct plasma membrane subdomains. This would represent a very different solution to generate more complex targeting to plasma membrane subdomains than the one found in animals. In the case of CS formation, CASPs and other CS proteins might be segregated into specific RabA-bearing secretory vesicle populations, that would then preferentially tether to PM domains with an EXO70A1 mark. In the endodermis EXO70A1 marks are distributed in centrally aligned microdomains through an as yet unknown mechanism.

## Materials and Methods

### Plant Material and growth conditions

For all experiments, *Arabidopsis thaliana* (ecotype Columbia) was used. Seeds were surface-sterilized, sown on plates contained half-strength Murashige and Skoog (MS) + 0.8% Agar (Roth) medium, stratified at 4°C and darkness for 2 days, and grown vertically in growth chambers at 21°C at constant light for 5 days, unless otherwise stated.

All plant materials used, either published or here obtained are listed in Table 1. The following available mutants were used *casp1-1* (SAIL_265_H05), *casp3-1* (SALK_011092), *casp5-1* (SALK_042116), *esb1-1* (Hosmani et al., 2013), *sgn3-3* (SALK_043282) and *myb36-2* (GK-543B11). Novel mutant alleles created in this study using CRISPR-Cas9 (Ursache et al., 2021a) (see protospacer sequences in Table 1) are *caspQ, caspQ sgn3, caspQ caspl-6ple*, using targeting constructs described below.

The following published constructs were introduced by floral dipping in caspQ: pCASP1::CASP1-GFP, pCASP2::CASP2-GFP, pCASP3::CASP3-GFP, pCASP4::CASP4:GFP, pCASP5::CASP5-GFP (Roppolo 2011 Nature), pCASP1::mCit-SYP122 (Pfister 2014 eLife), pESB1::ESB1-mCherry (Hosmani 2013 PNAS), pRBOHF:mCherry-RBOHF and pPER64::PER64-mCherry (Lee 2013 Cell); pCASP1::mVenus-SYP122 and pELTP::3xmCherry-SYP122 (Fujita 2020 EMBO J); pCASP1::CASP1-mCherry (Kalmbach et al., 2017) and pELTPxve::GELP72 (Ursache et al., 2021).

### Cloning and plasmid construction

The In-Fusion Advantage PCR Cloning Kit (Clontech) and Gateway Cloning Technology (Invitrogen) were used for generating lines described thereafter. All constructs were transformed by heat shock into Agrobacterium tumefaciens GV3101 strain and then transformed into plants by floral dipping.

CRISPR-Cas9 targeting of genes CASP1, CASP2, CASP4, SGN3, CASPL1-A1, B1, B2, C1, C2, D1 and D2 were generated as reported in (Ursache et al., 2021a). Briefly, on or two protospacer sequences were selected in online tool (Benchling). Forward and reverse primers containing protospacer sequences flanked by overhanging sites for each entry vector were annealed and ligated into entry clones. Alternating U6-26::gRNA and U3::gRNA containing vectors were assembled into intermediate vector at desired combinations by Golden-Gate reaction. Final vectors containing gRNA multiplexed ORFs, pUbi::Cas9 and FastRed selection were then generated by Gateway reaction. In Table 1, protospacer sequences are listed in *Primer* list, final multiplexing vectors in *Plasmid* list and the resulting homozygous plant alleles listed and described in *Plants* list.

The pCASPL*n*::CASPL*n*-mCitrine constructs were generated by Infusion as follows: CASPL promoter and gene sequences, without STOP codon, were amplified with overlapping primers to mCitrine and introduced by Infusion into destination vector containing FastRed cassette (see plasmid and primer lists in Table1).

To generate single double and triple CASP1-CASP5 combination constructs, CASP genes (promoter, gene and 3’UTR and terminator sequences) and GFP and/or mTurquoise tag were amplified by PCR with overlapping primers and cloned by Infusion into linearized gateway entry vectors, for any of 3 positions of Multi-site triple Gateway. Triple LR reactions were performed with different combinations of CASP1 to CASP5 genes, and in case of single and double CASP combinations, dummy entry vectors were used at remaining positions. All construct combinations and respective primers are listed in Table 1.

The pSCR::EXO70A1-mVenus construct was generated by LR from available entry vectors (Kalmbach et al., 2017).

For TurboID constructs, the sequences were amplified using primers listed in Table1, from vectors kindly provided by P. Moschou (Arora et al., 2020) for BP reaction to Gateway entry clones, and translational fusions were performed with Multi-site Gateway reactions.

The pCASP1::GFP-RabA*n* constructs were generated by triple Gateway. Briefly, Rab-GTPases were amplified and introduced by BP reaction in pDNR p2r-p3. Dominant negative mutations (N-I) were introduced by PCR site-directed mutagenesis in respective entry vectors pDONR p2r-p3. Final vectors were assembled by Multi-site Gateway reactions with pCASP1 promoter and GFP for N-terminal fusion, in Fast-Red selection vector.

### RNA extraction and qRT-PCR

For RNA extraction, seedlings were grown for 5 days on half strength MS on mesh to facilitate root cutting. Approximately 60 mg roots per biological replicate were cut and frozen in liquid nitrogen and stored at −80°C. Total RNA was extracted using a TRIzol-adapted ReliaPrep RNA Tissue Miniprep Kit (Promega). Reverse transcription was carried out with PrimeScript RT Master Mix (Takara). All steps were done as indicated in the manufacturer’s protocols. The qPCR was performed on an Applied Biosystems QuantStudio3 thermocycler using a MESA BLUE SYBR Green kit (Eurogentech). Transcripts are normalized to Clathrin adaptor complexes medium subunit family protein (AT4G24550) expression. All primer sets are indicated in Table 1.

### Confocal Laser Scanning Microscopy

Confocal laser-scanning microscopy images were obtained using either a Zeiss LSM 880 (with Zen 2.1 SP3 Black edition), Leica SP8-X (with LasX 3.5.6.21594) or Leica Stelaris 5 (LAS X (2020) version 4.1.23273) microscopes. The following excitation and detection windows were used: GFP, Fluorol yellow: 488 nm, 500–530 nm; mVenus, mCITRINE: 514 nm, 505–530 nm; Propidium iodide: 561 nm, 590–650 nm; Calcofluor White: 405 nm, 430–485 nm; Basic Fuchsin: 561 nm, 600–630 nm.

Propidium iodide penetration assay was performed as described previously (Alassimone et al., 2010). Briefly, seedlings were stained in PI (10µg/mL) dissolved in water for 5 min and mount in same solution for imaging, with above defined settings. Scoring of endodermal cell number was initiated from onset of elongation, (defined as endodermal cell length being more than two times than width in the median, longitudinal section) until PI could not penetrate into the stele.

Lignin and cell wall staining was performed in fixed and cleared seedlings, following ClearSee-adapted protocol (Fujita et al., 2020). Briefly, 5-day-old seedlings were fixed in 3 ml 1 × PBS containing 4% paraformaldehyde for 1 h at room temperature in 12-well plates and washed twice with 3 ml 1 × PBS. Following fixation, the seedlings were cleared in 3 ml ClearSee solution under gentle shaking. After overnight clearing, the solution was exchanged to new ClearSee solution containing 0.2% Fuchsin and 0.1% Calcofluor White for lignin and cell wall staining, respectively, and incubated overnight. Next day samples were rinsed with fresh ClearSee solution for 30 min with gentle shaking, 2-3 times, and at least once overnight before observation.

Suberin was stained with Methanol-based Fluorol Yellow protocol (Fujita et al., 2020). Briefly, 5-day-old seedlings were fixed and cleared in methanol for at least 3 days at 4°C, exchanging the methanol at least once. The cleared seedlings were transferred to a freshly prepared solution of Fluorol Yellow 088 (0.01%, in methanol) and incubated for 1 h. The stained seedlings were rinsed in fresh methanol and transferred to a freshly prepared solution of aniline blue (0.5%, in methanol) for counterstaining for 1 h. Finally, the seedlings were washed for 2–3 min in water before imaging. For *caspQ* pELTPxve>>GELP72 assay, seeds were sown directly in half-strength MS containing 10 µM beta-estradiol, including control *caspQ*.

For plasmolysis, seedlings were firstly stained in PI (10µg/mL) for 5 minutes, and directly mount in a 0.8M mannitol solution just before imaging.

### Image analysis: CS particle analysis and surface profiles

All images were processed and analyzed in FIJI. All images were acquired with Leica SP8 or Stellaris at the described endodermal cell number with fixed dimensions: 63x objective, 4x optical zoom, 1024 x 512 pixel, such that CS length was approximately equal among samples, *i.e.* 46 µm. At desired cell number, a region with most flat CS surface was chosen, and a Z-stack was acquired with as many stacks needed to capture whole strip in focus. For each image, stacks were projected with Maximum Intensity projection and processed as follows. For CS particle analysis, images were then analyzed with the “Analyze Particles” FIJI plugin as follows: a segmented line of 50 pt was drawn along the CS and straightened to make a ROI. The Threshold “InterModes” was applied to straighten strip, and plugin “Analyze Particles” run. Shown is *Particle number* per approximately 46µm of CS length; *Particle area* (µm^2^) and *CS area (%)* to Col-0 at cell 20.

For CS surface profiles, a segmented line of 20 pt width was drawn along the strip, and a profile was obtained for each channel. In Excel, each channel intensities were normalized to average intensity and plotted in GraphPad.

### Classical TEM analysis

Plants were fixed in glutaraldehyde solution (EMS, Hat-field, PA) 2.5% in phosphate buffer (PB 0.1 M [pH 7.4]) for 1 h at RT and postfixed in a fresh mixture of osmium tetroxide 1% (EMS, Hatfield, PA) with 1.5% of potassium ferrocyanide (Sigma, St. Louis, MO) in PB buffer for 1 h at RT. The samples were then washed twice in distilled water and dehydrated in ethanol solution (Sigma, St Louis, MO, US) at graded concentrations (30%–40 min; 50%–40 min; 70%–40 min; 100%–2x 1 h). This was followed by infiltration in Spurr resin (EMS, Hatfield, PA, US) at graded concentrations (Spurr 33% in ethanol - 4 h; Spurr 66% in ethanol - 4 h; Spurr 100%–2x 8 h) and finally polymerized for 48 h at 60 °C in an oven. Ultrathin sections of 50 nm thick were cut transversally at 1.8mm from the root tip using a Leica Ultracut (Leica Mikrosysteme GmbH, Vienna, Austria), picked up on a copper slot grid 2 x 1 mm (EMS, Hatfield, PA, US) coated with a polystyrene film (Sigma, St Louis, MO, US). For sequential cut, the roots were cut every 0.1mm starting from 0.9mm from the root tip. Sections were post-stained with uranyl acetate (Sigma, St Louis, MO, US) 4% in H2O for 10 min, rinsed several times with H2O followed by Reynolds lead citrate in H2O (Sigma, St Louis, MO, US) for 10 min and rinsed several times with H2O. Micrographs were taken with a transmission electron microscope Philips CM100 (Thermo Fisher Scientific, Waltham, MA USA) at an acceleration voltage of 80 kV with a TVIPS TemCamF416 digital camera (TVIPS GmbH, Gauting, Germany) using the software EM-MENU 4.0 (TVIPS GmbH, Gauting, Germany). Panoramic alignments were performed with the software IMOD (Kremer et al., 1996).

### High Pressure freezing and cryo-substitution

For the High Pressure Freezing, pieces of root 5 mm long were cut from tip, and then placed in an aluminum planchet of 6 mm in diameter with a cavity of 0.1 mm (Art.610, Wohlwend GmbH, Sennwald, Switzerland) filled with 15% Dextran in 2-morpholinoethanesulfonic acid buffer (MES 50 mM, [pH 5.7]) covered with a tap planchet (Art.611, Wohl-wend GmbH, Sennwald, Switzerland) and directly high pressure freezed using a High Pressure Freezing Machine HPF Compact 02 (Wohlwend GmbH, Sennwald, Switzerland). The samples were then chemically fixed, dehydrated and infiltrated with resin at cold temperature using the Leica AFS2 freeze substitution machine (Leica Mikrosysteme GmbH, Vienna, Austria) with the following protocol: Dehydration and fixation in a solution containing a mixture of osmium tetroxide 0.5% (EMS, Hatfield, PA) with glutaraldehyde 0.5% (EMS, Hatfield, PA) with uranyl acetate 0.1% (Sigma, St. Louis, MO) in acetone (Sigma, St Louis, MO, US) at graded temperature (−90 °C for 30 h; from −90 °C to −60 °C in 6 h; −60 °C for 10 h; from −60 °C to −30 °C in 6 h; −30 °C for 10 h; from −30 °C to 0° in 6 h) This was followed by washing in acetone and then infiltration in Spurr resin (EMS, Hatfield, PA, US) at graded concentration and temperature (30% for 10 h from 0 °C to 20°C; 66% for 10h at 20°C; 100% twice for 10h at 20°C) and finally polymerized for 48 h at 60 °C in an oven.

### TEM tomography and 3D reconstruction

For electron tomography, semi-thin sections of 250 nm thickness were cut transversally to the root using a Leica Ultracut (Leica Mikrosysteme GmbH, Vienna, Austria) and then, picked up on 75 square mesh copper grids (EMS, Hatfield, PA, US). Sections were post-stained on both sides with uranyl acetate (Sigma, St Louis, MO, US) 2% in H2O for 10 min and rinsed several times with H2O. Protein A Gold 10 nm beads (Aurion, Wageningen, The Netherlands) were applied as fiducials on both sides of the sections and the grids were placed on a dual axis tomography holder (Model 2040, Fischione Instruments). The area of interest was taken with a transmission electron microscope JEOL JEM-2100Plus (JEOL Ltd., Akishima, Tokyo, Japan) at an acceleration voltage of 200 kV with a TVIPS TemCamXF416 digital camera (TVIPS GmbH, Gauting, Germany) using the SerialEM software (Mastronarde, D.N., 2005). Micrographs were taken as single tilt series over a range of −60° to +60° using SerialEM at tilt angle increment of 1°. Tomogram reconstruction, segmentation and model visualization were done with IMOD software (Kremer et al., 1996).

### Lignin staining with permanganate potassium (KMnO_4_) using TEM

Visualization of lignin deposition around Casparian strip was done using permanganate potassium (KMnO_4_) staining (Hepler et al., 1970). The sections were post-stained using 1% of KMnO_4_ in H_2_O (Sigma, St Louis, MO, US) for 45 min and rinsed several times with H_2_O. Micrographs were taken with a transmission electron microscope FEI CM100 (FEI, Eindhoven, The Netherlands) at an acceleration voltage of 80kV with a TVIPS TemCamF416 digital camera (TVIPS GmbH, Gauting, Germany) using the software EM-MENU 4.0 (TVIPS GmbH, Gauting, Germany). Panoramic were aligned with the software IMOD .

### Detection of H_2_O_2_ production in situ using transmission electron microscopy

Visualization of H_2_O_2_ production around Casparian strip and *caspQ* dots was done by cerium chloride method as described previously (Bestwick et al., 1997; Lee et al., 2013). Five day old *Arabidopsis* seedlings were incubated in 50 mM MOPS pH7.2 containing 10 mM CeCl_3_ for 30 min. After incubation with CeCl_3_, seedlings were washed twice in MOPS buffer for 5 min and fixed in glutaraldehyde solution (EMS, Hatfield, PA) 2.5% in 100 mM phosphate buffer (pH 7.4) for 1 h at room temperature. Then, they were post fixed in osmium tetroxide 1% (EMS) with 1.5% of potassium ferrocyanide (Sigma, St. Louis, MO) in phosphate buffer for 1 h at room temperature. Following that, the plants were rinsed twice in distilled water and dehydrated in ethanol solution (Sigma) at gradient concentrations (30% 40 min; 50% 40 min; 70% 40 min; two times (100% 1 h). This was followed by infiltration in Spurr resin (EMS) at gradient concentrations [Spurr 33% in ethanol, 4 h; Spurr 66% in ethanol, 4 h; Spurr two times (100% 8 h)] and finally polymerized for 48 h at 60°C in an oven. Ultrathin sections 50 nm thick were cut transversally at 1.8 ± 0.1 mm from the root tip, on a Leica Ultracut (Leica Microsystems GmbH, Vienna, Austria) and picked up on a copper slot grid 2 × 1 mm (EMS) coated with a polystyrene film (Sigma). Micrographs were taken with a transmission electron microscope FEI CM100 (FEI, Eindhoven, The Netherlands) at an acceleration voltage of 80 kV with a TVIPS TemCamF416 digital camera (TVIPS GmbH, Gauting, Germany) using the software EM MENU 4.0 (TVIPS GmbH, Gauting, Germany).

### Biotin labelling, Protein extraction and Western Blotting

#### Biotin labelling

For each sample, 160 mg of surface-sterilized seeds were imbibed in 0.2% agar and sown in described media containing mesh using 1mL pipette along two lines per square plate. Ater 2 days stratification in darkness at 4°C, seedlings were grown for 6 days at 22°C continuous light, before transferring to 50 µM biotin containing half-strength MS media, under sterile conditions using the mesh to transfer whole seedling population of each plate. Seedlings were grown for another 24h in biotin containing plates at 28°C continuous light. One biological replicate, from each genotyped, comprised 10 plates.

#### Protein Extraction

Roots (100-120 mg per plate) were collected with razor blade, weighted and immediately frozen in 2mL eppis containing a 5mm metal bead in liquid nitrogen. One biological replicate from 10 plates, yielded app. 1g of root material. Roots were ground using Tissue Lyzer II (Qiagen) with 2 cycles, 30sec, maximum speed, taking care to maintain material frozen, using pre-cooled racks at −80°C and freezing samples in liquid-N_2_ between cycles. Protein extraction and biotinylated-protein purification was done adapting available protocols (Hung et al., 2016; Mair et al., 2019) as follows: Ground roots were resuspended with Volume (2x weight) of RIPA buffer containing protease inhibitors (50 mM Tris, 150 mM NaCl, 0.1% (wt/vol) SDS, 0.5% (wt/vol) sodium deoxycholate and 1% (vol/vol) Triton X-100 in Millipore water, adjusted pH to 7.5 with HCl, 1mM PMSF and 2.5x PIC-Complete (Roche) in cold by short vortexing and spin down at 16’000 g for 10min at 4° C. For checking success of biotin feeding by western blot (*input*), 10 µL of extract were boiled in 4x Laemmli buffer supplemented with 20 mM DTT and 2 mM biotin at 95°C for 5 min, and western blots performed as described below.

Supernatants were collected and dialyzed to remove free biotin with PD-10 columns (GE Healthcare) following manufacturer’s instructions. Dialysed extracts (app. 3mL) were then incubated with 200µL Dynabeads MyOne Streptavidin C1 (Invitrogen) overnight. Next day, beads were separated from protein extract using a magnetic rack, and washed with following sequence of buffers: 2x cold RIPA buffer, 1x cold KCl 1 M, 1x with cold, 1x cold 100 mM Na_2_CO_3_, 1x room temperature 2M Urea in 10 mM Tris pH 8, and 2x cold RIPA buffer. For checking success of purification, 2% of the beads were boiled in 50 ml 4x Laemmli buffer supplemented with 20 mM DTT and 2 mM biotin at 95°C for 5 min for immunoblots (*purified*). The rest of the beads was spun down to remove the remaining wash buffer, added 1x PBS and stored at −80°C until further processing.

Western Blots of *input* and *purified* samples, proteins were separated in 4-12% gradient SDS-PAGE (Thermoscientific), and semi-dry transferred to Nitrocellulose membranes (Thermoscientific Pierce G2). For anti-GFP, membranes were blocked in PBS 0.5% Tween-20 5% Milk overnight, incubated 1h in anti-GFP (rabbit polyclonal, 1:3000, Invitrogen), washed in 3x 10min PBS Tween 0.5%, incubated in anti-rabbit-HRP (1:5000, Promega) and washed in 3x 10min PBS Tween 0.5% before detection. For streptavidin-HRP, membranes were blocked in TBS 0.1% Tween-20 5% Milk overnight, incubated 1h in streptavidin-HRP (1:2000, ThermoScientific), washed in 3x 10min TBS 0.1% Tween-20, 1x 10min TBS and 1x 10min destilled water before detection. Chemiluminescence was detected using SuperSignal West Pico kit (ThermoScientific).

### Mass-Spectrometry

#### Sample preparation

Protein digestion protocol on beads was adapted from (Han et al., 2017). Magnetic beads were washed with 500 ul of 2M urea / 50 mM TEAB (pH 7.8) and resuspended in 100 ul of the same solution suplemented with 1 mM DTT and 0.5 ug trypsin. After 1 h of incubation at 25°C, the supernatant was collected, beads washed with 100 ul of 2 M urea / 50 mM TEAB and this second supernatant pooled with the first one. After addition of 4 mM DTT (final concentration), the samples were reduced 30 min at 25°C, and then alkylated with 20 mM chloroacetamide (final concentration) during 45 min at 25°C in the dark. After addition of 0.5 ug of trypsin, the samples were digested overnight at 25°C. The solutions were then acidified with 10 ul of 20% trifluoroacetic acid, desalted on a Sep-Pak tC18 Waters plate and eluted with 150 ul of 80% acetonitrile / 0.1% formic acid.

#### Mass spectrometry analyses

Tryptic peptides fractions were dried and resuspended in 0.05% trifluoroacetic acid, 2% (v/v) acetonitrile, for mass spectrometry analyses. Tryptic peptide mixtures were injected on an Ultimate RSLC 3000 nanoHPLC system (Dionex, Sunnyvale, CA, USA) interfaced to an Orbitrap Fusion Tribrid mass spectrometer (Thermo Scientific, Bremen, Germany). Peptides were loaded onto a trapping microcolumn Acclaim PepMap100 C18 (20 mm x 100 μm ID, 5 μm, 100Å, Thermo Scientific) before separation on a reversed-phase custom packed nanocolumn (75 μm ID × 40 cm, 1.8 μm particles, Reprosil Pur, Dr. Maisch). A flowrate of 0.25 μl/min was used with a gradient from 4 to 76% acetonitrile in 0.1% formic acid (total time: 140 min). Full survey scans were performed at a 120’000 resolution, and a top speed precursor selection strategy was applied to maximize acquisition of peptide tandem MS spectra with a maximum cycle time of 0.6s. HCD fragmentation mode was used at a normalized collision energy of 32%, with a precursor isolation window of 1.6 m/z, and MS/MS spectra were acquired in the ion trap. Peptides selected for MS/MS were excluded from further fragmentation during 60s.

#### Data analysis

Tandem MS data were processed by the MaxQuant software (version 1.6.3.4) (Cox and Mann, 2008) incorporating the Andromeda search engine (Cox et al., 2011). An *Arabidopsis thaliana* reference proteome database from UniProt (November 2019 version, 39’362 sequences), supplemented with sequences of common contaminants, was used. Trypsin (cleavage at K,R) was specified as the enzyme definition, allowing 2 missed cleavages. Carbamidomethylation of cysteine was specified as a fixed modification, N-terminal acetylation of protein and oxidation of methionine as variable modifications. All identifications were filtered at 1% FDR at both the peptide and protein levels with default MaxQuant parameters. For protein quantitation the iBAQ values (Schwanhäusser et al., 2011) were used. MaxQuant data were further processed with Perseus software (version 1.6.14.0) (Tyanova et al., 2016) for the filtering, log2-transformation and normalization of values, data imputation, statistical analyses and GO annotations. The mass spectrometry proteomics data have been deposited to the ProteomeXchange Consortium via the PRIDE partner repository. GO annotation enrichment test was performed based on (Cox and Mann, 2012). Sub-cellular localizations were queried in SUBA4 database (Hooper et al., 2017) following consensus algorithm.

#### Statistical analysis

All experiments were analysed in Excel, and statistical tests and graphs done in Prism GraphPad 9. Unless otherwise stated, data were analysed by One-Way ANOVA, and multiple comparisons against a selected control (wild-type or *caspQ*) were performed with Dunnet’s test or for all combinations with Tukey test, and significance level selected at *p*<0.05.

## Figure Legends

***Table 1: List of materials plant, plasmid and primers***

List of materials used in this study, generated here or elsewhere: *Plants*, with mutation alleles, transgene and resistances; *Plasmids*, used for transgenics and CRISPR-Cas9 with respective and *Primers*: including protospacer sequences used for CRISPR-Cas9; and primers used for qPCR and cloning of marker lines.

***Table 2: LC-MS/MS identified proteins in endodermal TurboID experiment. Sheet1)***

List of all proteins identified in biological replicates of TurboID samples. Protein abundances are given in log2 iBAQ (‘intensity-based absolute quantification’, see main text), all parameters of paired *t*-test (T-test statistic, Mean difference, p-value and *q*-value (adjusted p-value with Bonferroni correction)) and LC/MS-MS peptide parameters per protein are also given. For statistical purposes, proteins with non-detected values in one biological replicate were imputed values using normal distribution method of MaxQuant, Perseus software (version 1.6.14.0). No paired *t-*tests were performed if proteins were absent in all replicates of both compared samples (*p-*value N/A). For statistics, *q*-value < 0.05 was considered. **Sheet2)** List of 332 proteins significantly enriched in CASP1-tID samples in all pairwise combinations.

***Movie 1: Tomogram of caspQ foci after chemically fixation.***

Animated tomograms of *caspQ* foci after chemically fixation from semi-thin sections of 250 nm thickness. Micrographs were taken as single tilt series over a range of −60° to +60° using SerialEM at tilt angle increment of 1°. Tomogram reconstruction, segmentation and model visualization were done with IMOD software (Kremer et al., 1996). In 3D, chemical fixation induced-plasmolysis, allows visualization of PM detachment (blue) throughout the structure, presence of extracellular vesicular bodies (yellow) and flat lamellar structure (dark yellow) interspersed within the lignified cell wall (green) of *caspQ* focci.

***Movie 2: Tomogram of caspQ foci after cryo-fixation.***

Animated tomograms of *caspQ* foci after high-pressure freeze-substitution from semi-thin sections of 250 nm thickness. Micrographs were taken and processed as in movie 1. High-pressure freezing allows the reconstruction of presumably turgid cell, where membrane is compressed against apoplastic space, but plasma membrane and cell wall are still separated by the presence of extracellular vesicular bodies, as in the chemically fixed sample. Thin transversal aligned lamellae inside the *caspQ* foci, are also detected by this method.

## Acknowledgments

We thank Tonni Grube Andersen for sharing unpublished TRAP RNAseq data and Panagiotis M. Mouschou for sharing the unpublished TurboID construct. We thank Patrice Waridel and Manfredo Quadroni from Protein Analysis Facility of University of Lausanne, for support in the design, sample preparation and data analysis of TurboID experiment. This work was supported by EMBO long-term fellowship ALTF 987-2017 to I.B., an AgreenSkills+ fellowship (grant agreement no. FP7-609398) to K.H. and two consecutive Swiss National Science Foundation (SNSF) grants (project numbers 176399 and 156261) to N.G.

---

**Supplementary Figure 1.1:**
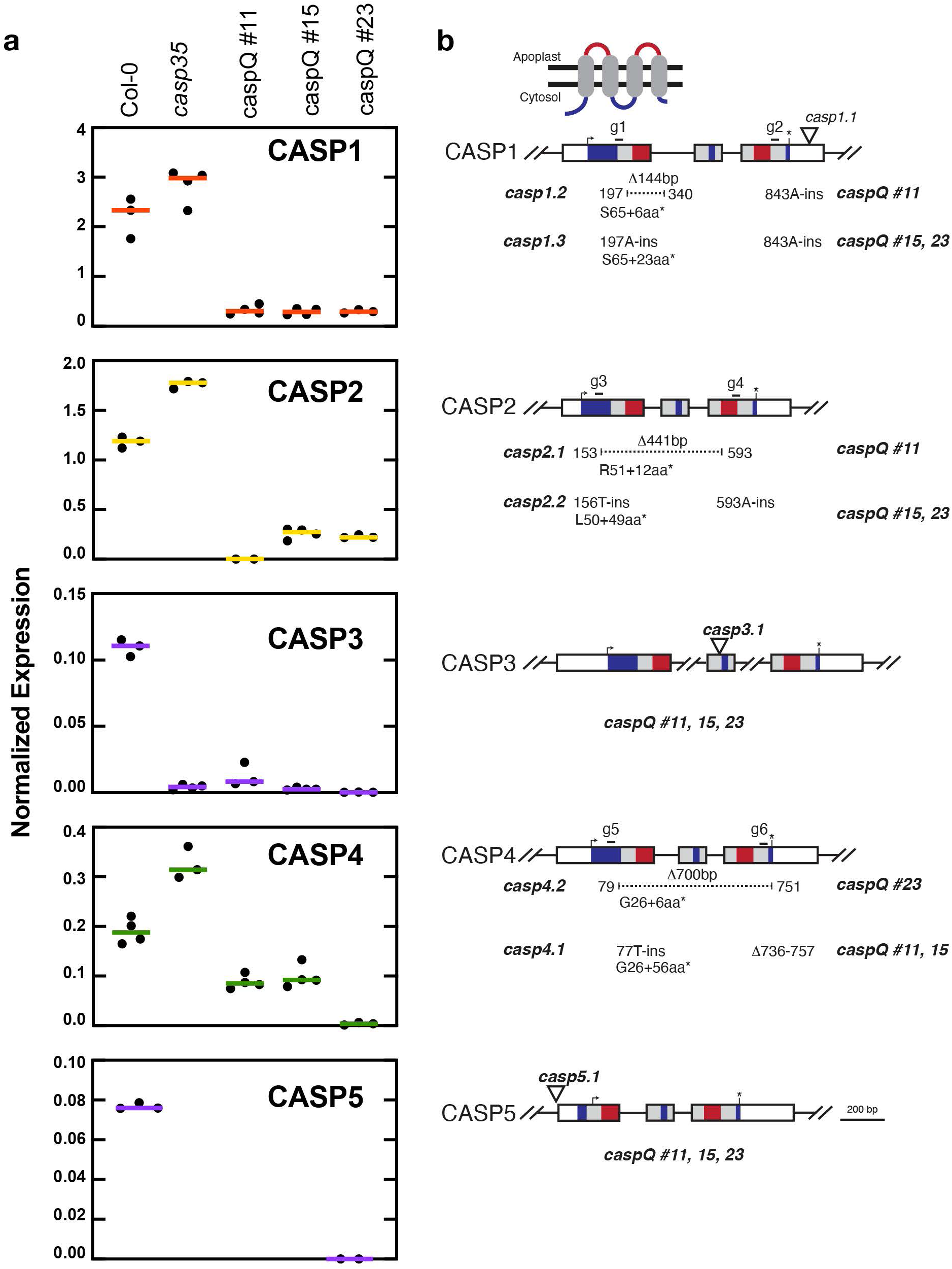
CASP expression and CRISPR-Cas9 targeting strategy in caspQ. **a.** Normalized gene expression of CASP1-CASP5 genes in wild-type Col-0, T-DNA mutant *casp35* and CRISPR-Cas9 *caspQ* alleles. Expression normalized to *Clathrin adaptor complexes medium subunit family protein* (AT4G24550). **b.** CRISPR-Cas9 gene targeting of *CASP1*, *CASP2* and *CASP4* and predicted early STOP codon alleles in *casp35* T-DNA mutant background. White boxes indicate 5’ and 3’UTR, coloured boxes predicted protein domains (blue, intracellular loops; gray, transmembrane domains; red, extracellular domains); top-right arrow, START; *, STOP codon; *g1-g6*, guide RNAs; position. Δ, deletions; ins, insertions; aa, amino-acid

**Supplementary Figure 1.2:**
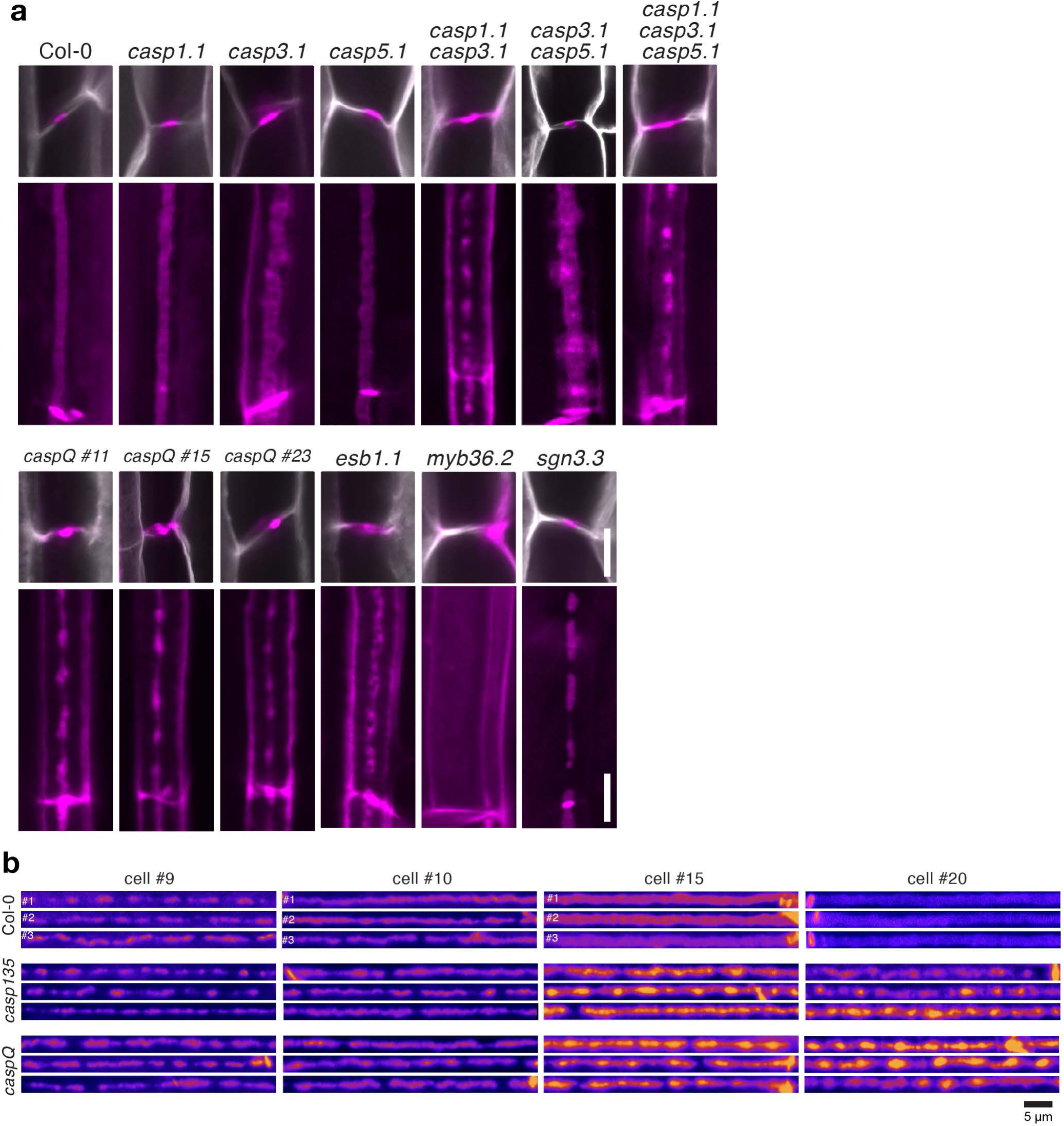
Representative pictures of lignification in mid and surface views. **a.** Mid (upper) and surface (lower panel) views of lignin (magenta, Basic Fuchsin) and cell wall (gray, calcofluor white) staining at endodermal cell number 20 of additional mutants analyzed in Figure 1. Pictures of same genotypes are identical to main figure. **b.** Representative pictures of CS surface view from three individuals at endodermal cell numbers 9, 10, 15 and 20 from Col-0, *casp135* and *caspQ.* Scale bars 5 µm.

**Supplementary Figure 1.3:**
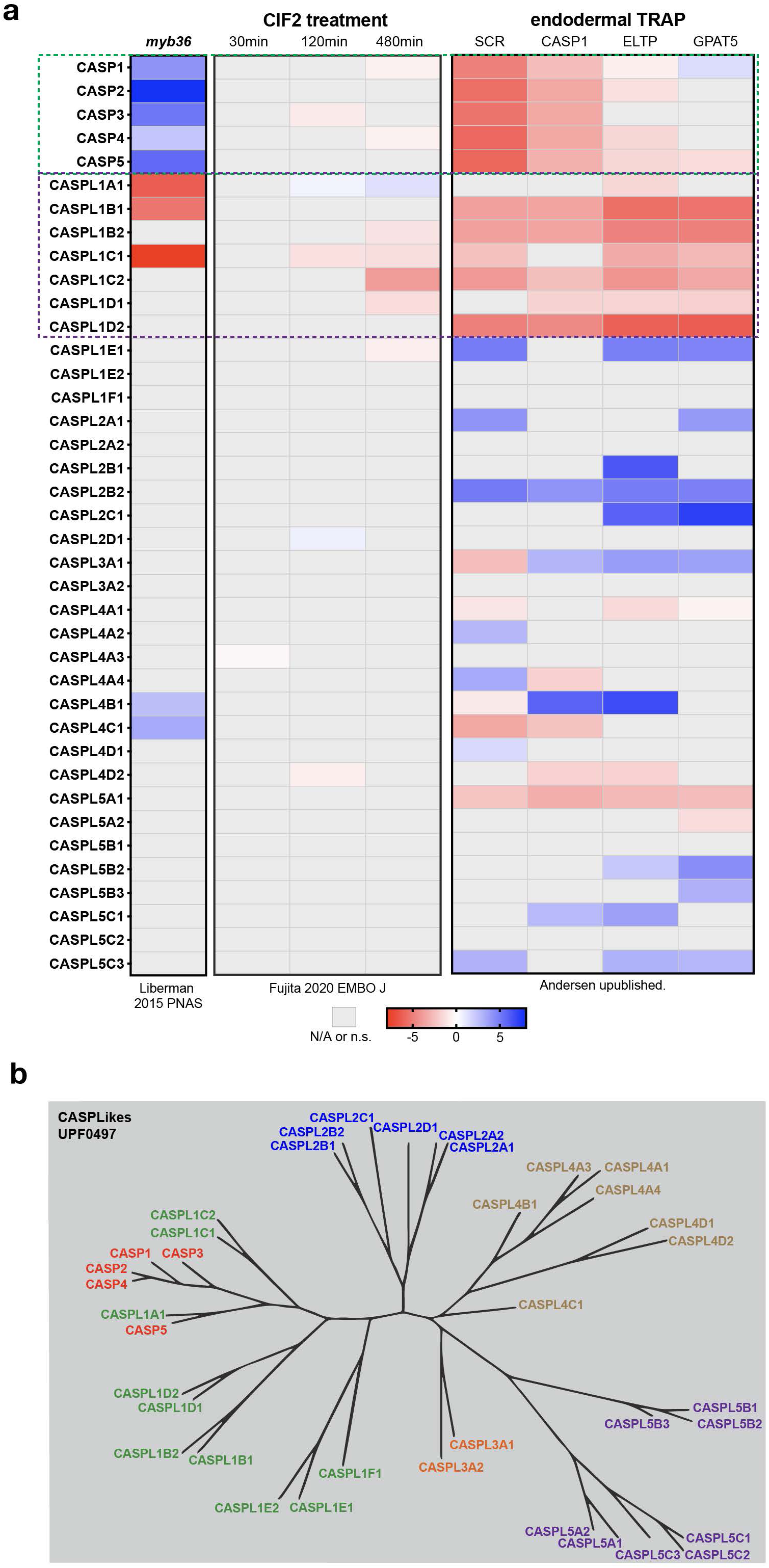
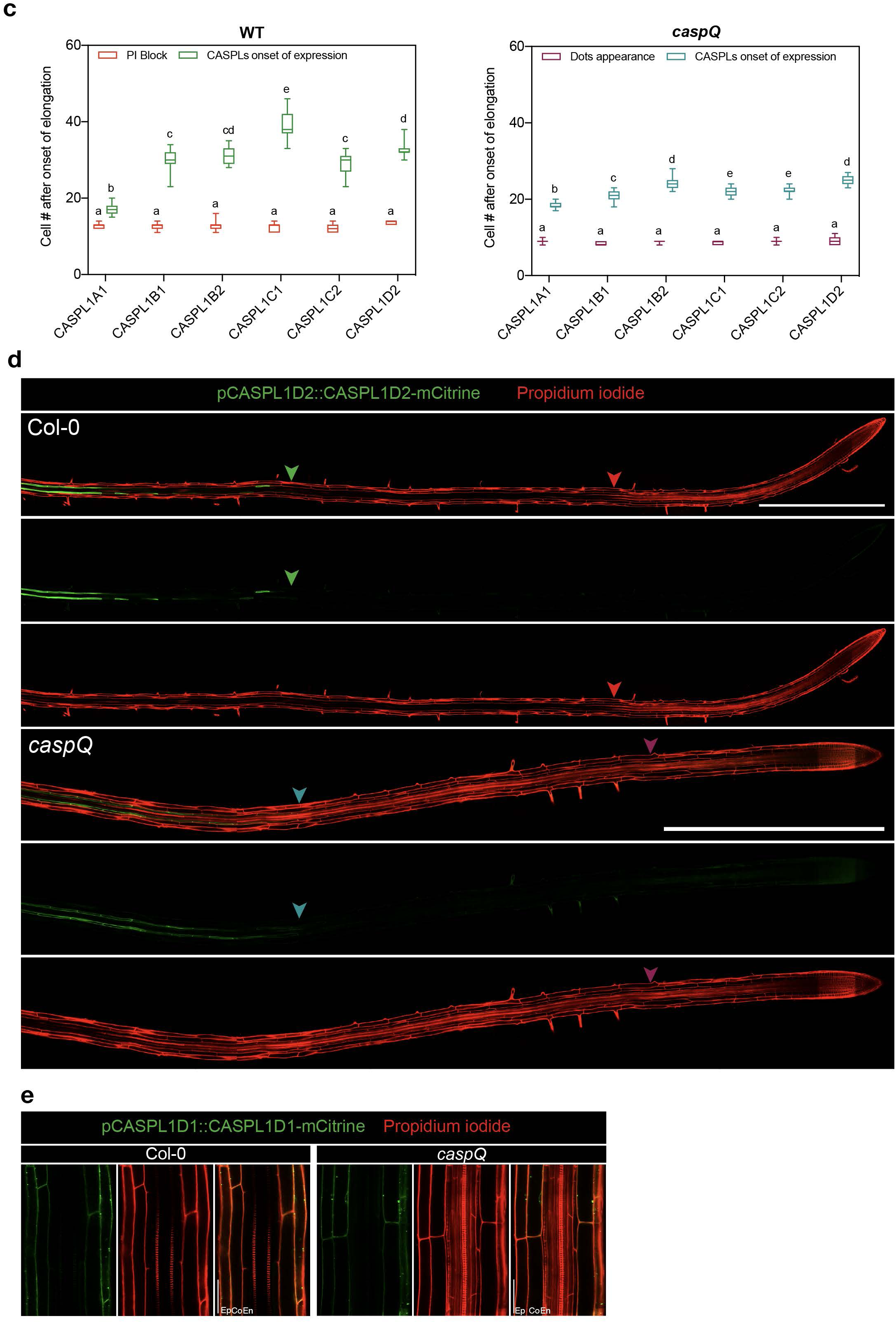
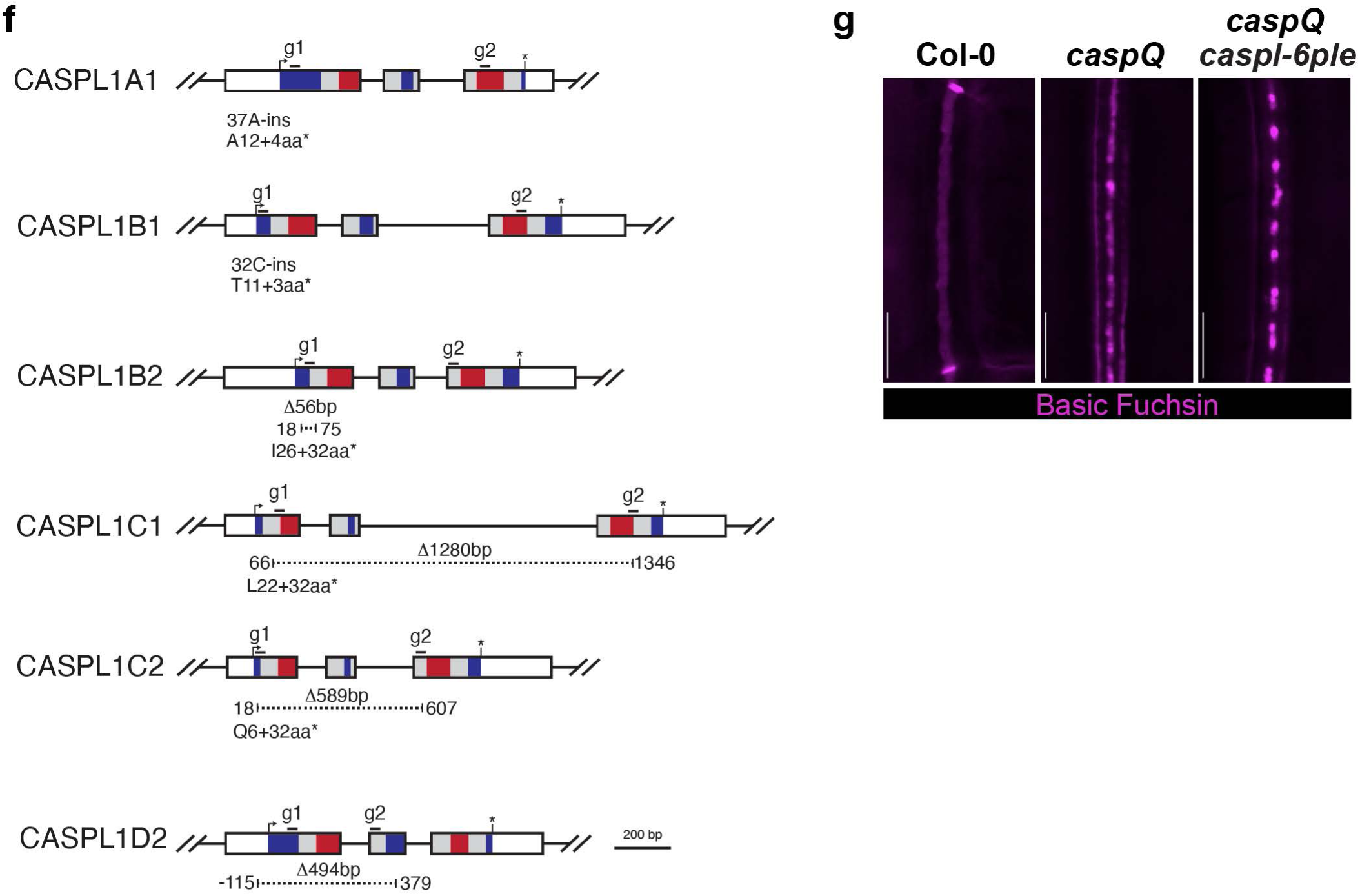
Other CASP-LIKES are not involved in CS formation. **a.** Comparative expression data of CASP-LIKE genes in endodermal and/or CS-relevant RNA-seq experiments: *myb36* mutant differential expression (Liberman et al., 2015); CIF2 treatment (Fujita et al., 2020) and endodermal TRAP-Seq (Andersen et al., *unpublished*) **b.** CASPL phylogenetic tree, adapted from (Roppolo et al., 2014). **c-d.** Analysis of spatial expression of translational fusions *pCASPLx::CASPLx-mCitrine* (*CASPL1A1, CASPL1B1, CASPL1B2, CASPL1C1, CASPL1C2* and *CASPL1D2*) in Col-0 and *caspQ* using PI-block (in Col-0) or PI-stained foci (in *caspQ*) as reference for endodermal differentiation. Different letter represent significant differences (p<0.05) by ANOVA and multiple comparisons Tukey test. **d.** Representative root pictures as taken for analysis in (c) for *pCASPL1D2::CASPL1D2-mCitrine*. Scale bar 1 mm. **e.** Representative pictures for pCASPL1D1::CASPL1D1-mCitrine at cell 20 after onset of elongation. Scale bar 50µm. **f.** CRISPR-Cas9 gene targeting of *CASPL1A1, CASPL1B1, CASPL1B2, CASPL1C1, CASPL1C2, CASPL1D1* and *CASPL1D2* in the *caspQ* background (*caspQ caspl-7ple*). White boxes 5’ and 3’UTR, coloured boxes predicted protein domains (blue, intracellular loops; gray, transmembrane domains; red, extracellular domains); top-right arrow, START; *, STOP codon; g1-g6, guide RNAs; Δ, deletions; ins, insertions; aa, amino-acid position. **g.** CS surface views stained with Basic Fuchsin (lignin, magenta) of Col-0, *caspQ* and *caspQ caspl-7ple*. Scale bar 10µm.

**Supplementary Figure 2.1:**
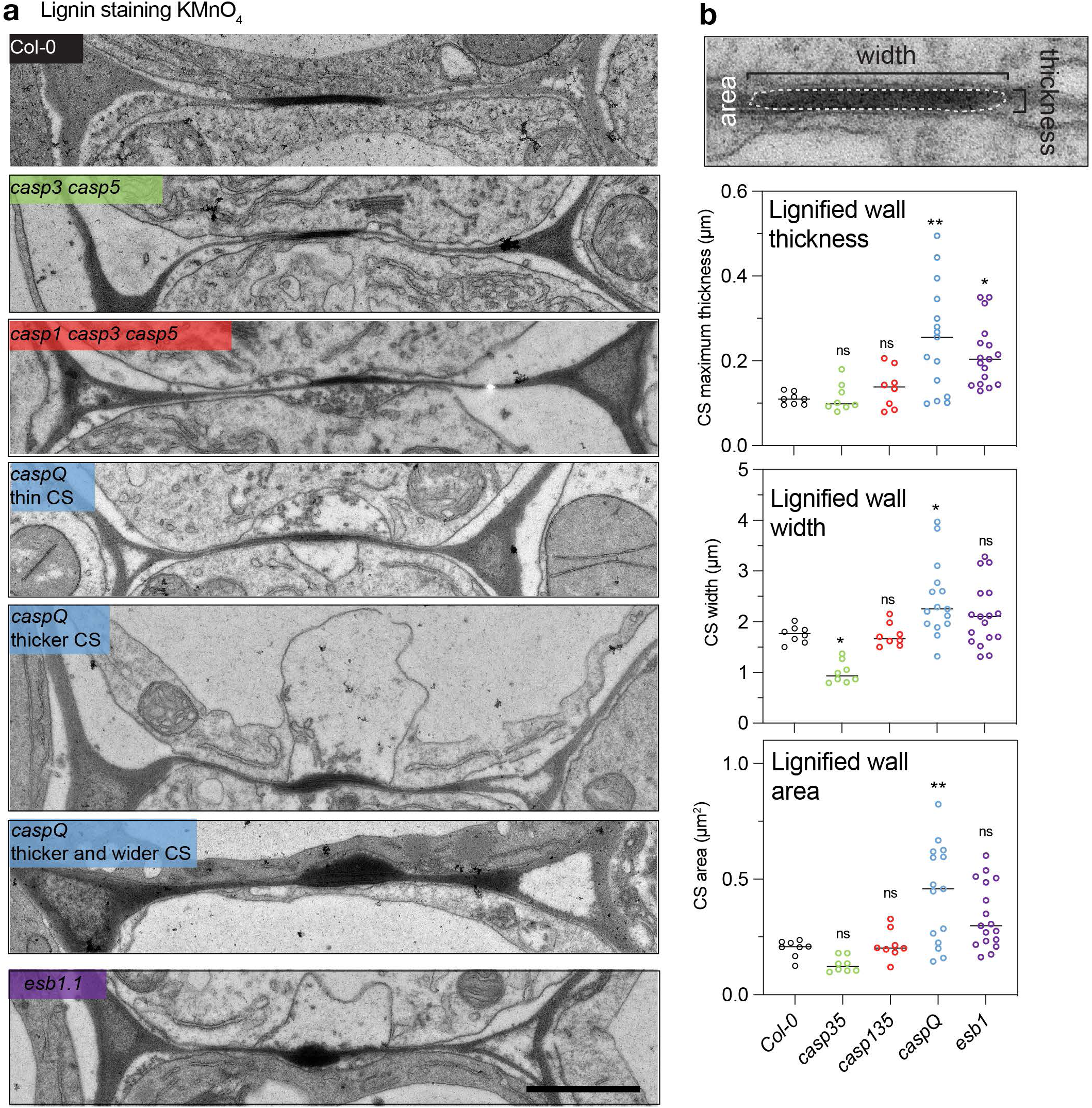
Representative electron-micrographs of all genotypes analyzed. **a.** Representative electron micrographs with KMnO_4_ lignin staining of endodermal cell-cell junctions in transversal cuts at 1.8 mm from root tip from wild-type, *casp35, casp135, caspQ* and *esb1*. For *caspQ,* three distinctive appearances are shown. **b**. Parameter quantification on central-aligned lignified walls stained by KMnO_4_: thickness, width and area. Graphs show measurements of n ≈ 16 cell-cell junctions from 2 individuals; mean (line) and significant differences to wild-type by ANOVA, Dunnett’s test *p*-value < 0.05. Scale bars 10 µm.

**Supplementary Figure 2.2:**
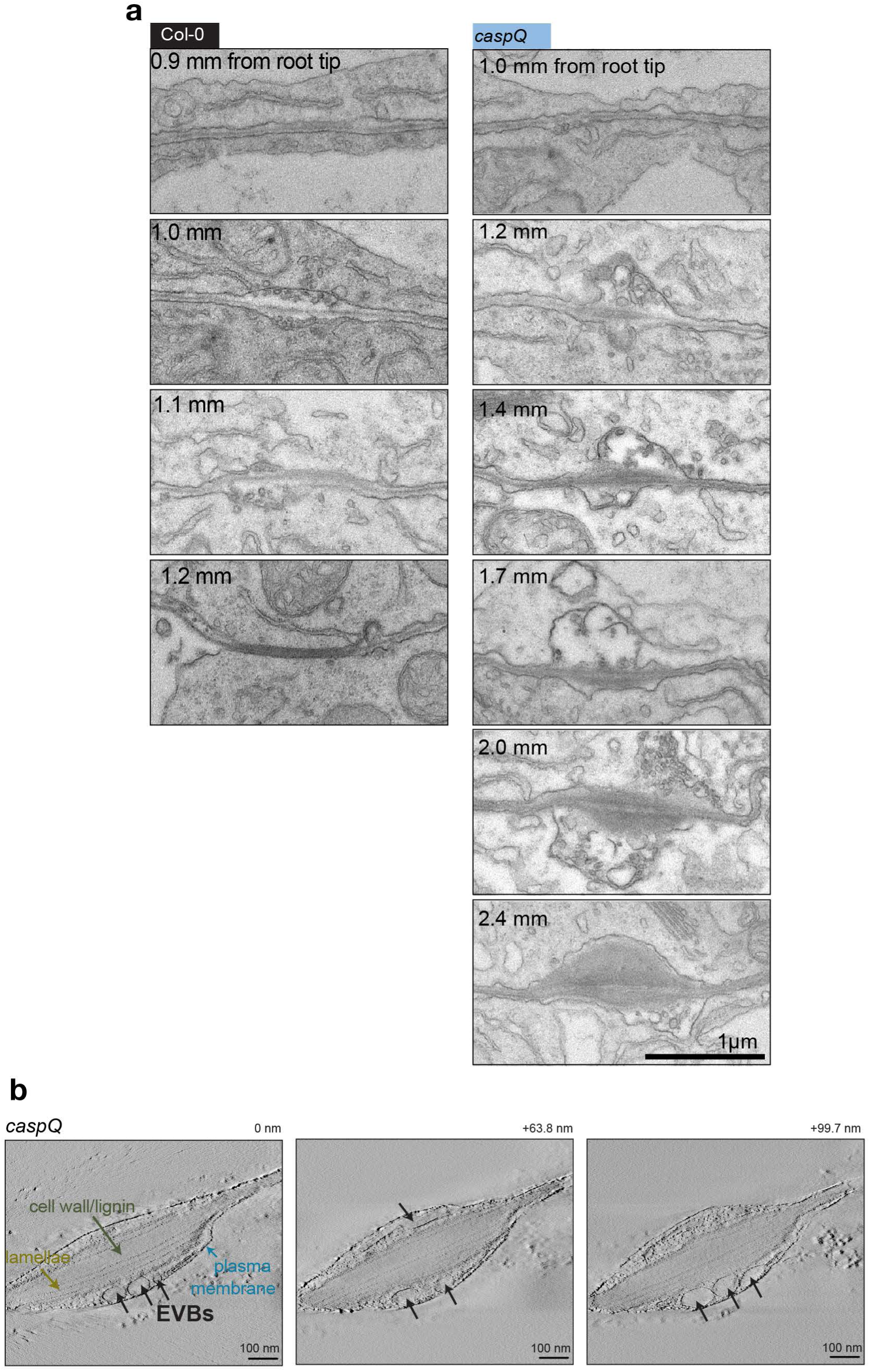
Series of electron-micrographs from manual series and tomograms. **a.** Representative electron micrographs as in Figure 2d of Col-0 and *caspQ* serial cuts along same seedling. Scale bar 1 µm. **b.** Series of three optical sections from the tomogram of high-pressure freezing samples from a lignified foci of *caspQ* mutant. Scale bar 0.1 µm.

**Supplementary Figure 2.3:**
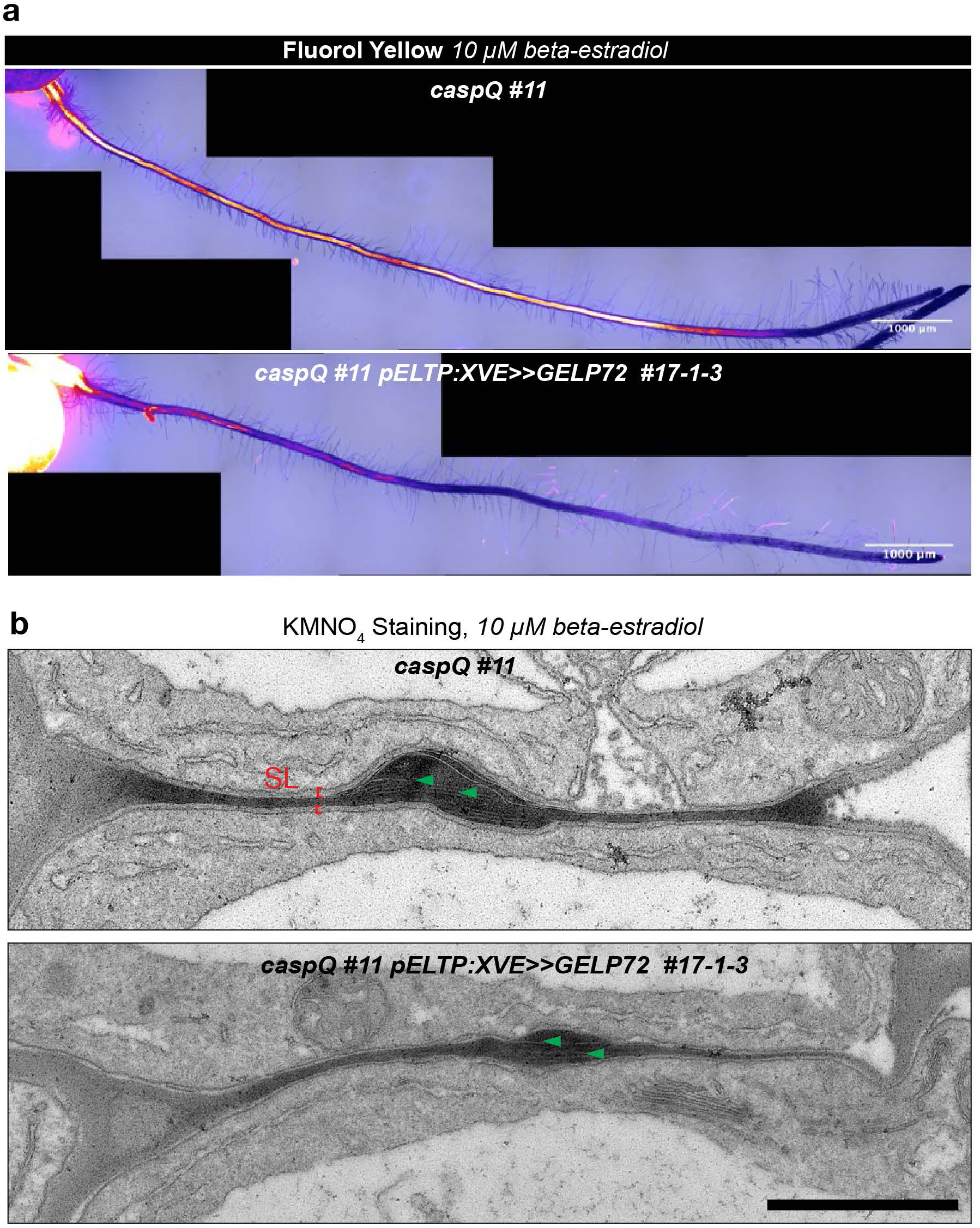
Lamellae inside caspQ lignin foci are not removed by suberin-degrading enzyme GELP72. **a-b.** Fluorol Yellow suberin staining **(a)** and Electron micrographs **(b)** of 5d old seedlings grown in 10µM β-estradiol of *caspQ* and *caspQ* pELTP:XVE>>GELP72 (pRU182) #17-1-3. Scale bars 1 mm (a) and 1 µm (b)

**Supplementary Figure 3.1:**
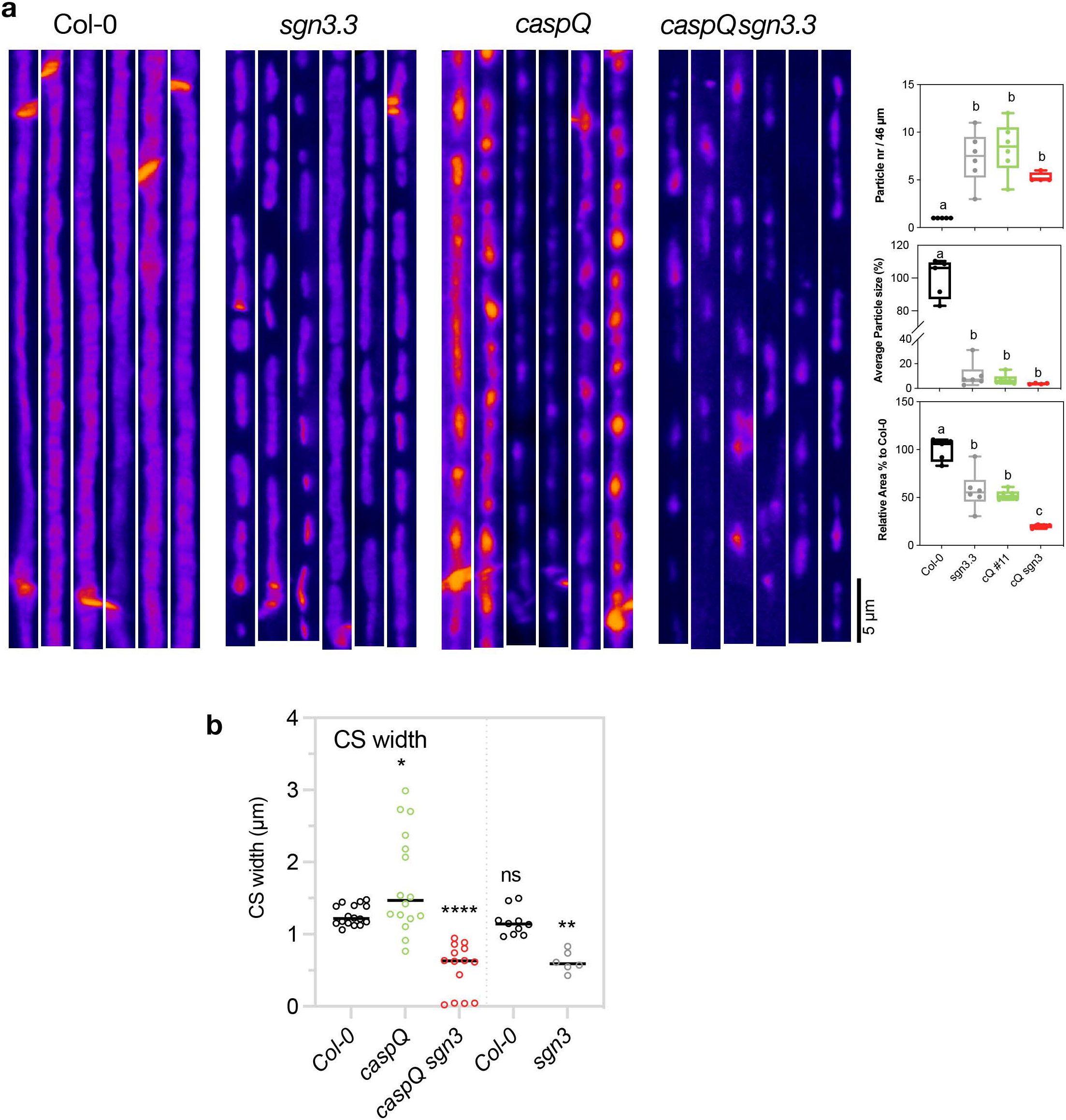
caspQ sgn3 lignification is strongly reduced. **a.** Pictures of CS surfaces used for particle analysis FIJI-plugin and respective derived parameters from Col-0, *sgn3*, *caspQ* and *caspQ sgn3*, as shown in Fig. 3b. **b.** Quantification CS width from EM pictures of *Col-0*, *sgn3*, *caspQ* and *caspQ sgn3* as shown in Fig.3c-d. Scale bar 5 µm.

**Supplementary Figure 3.2:**
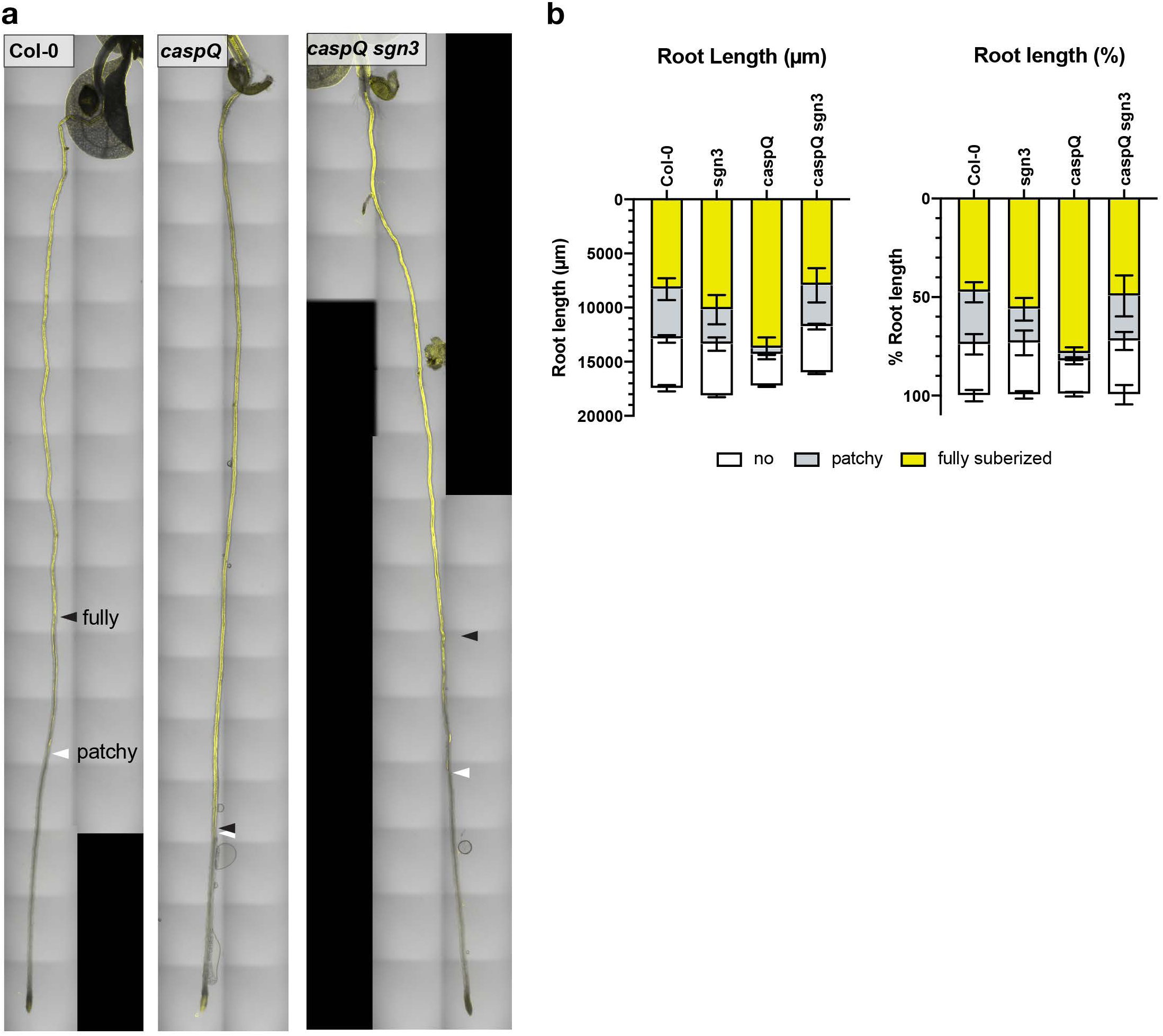
caspQ ectopic suberin is SGN3-dependent. **a-b.** Fluorol Yellow suberin staining in *Col-0*, *sgn3*, *caspQ* and *caspQ sgn3* 5d old-seedlings. Confocal pictures **(a)** and quantification of endodermal suberin patterns **(b):** no suberin (white), patchy (gray) and fully suberized (yellow) as function of absolute (left) and relative (%, right) root length to *Col-0*.

**Supplementary Figure 4.1:**
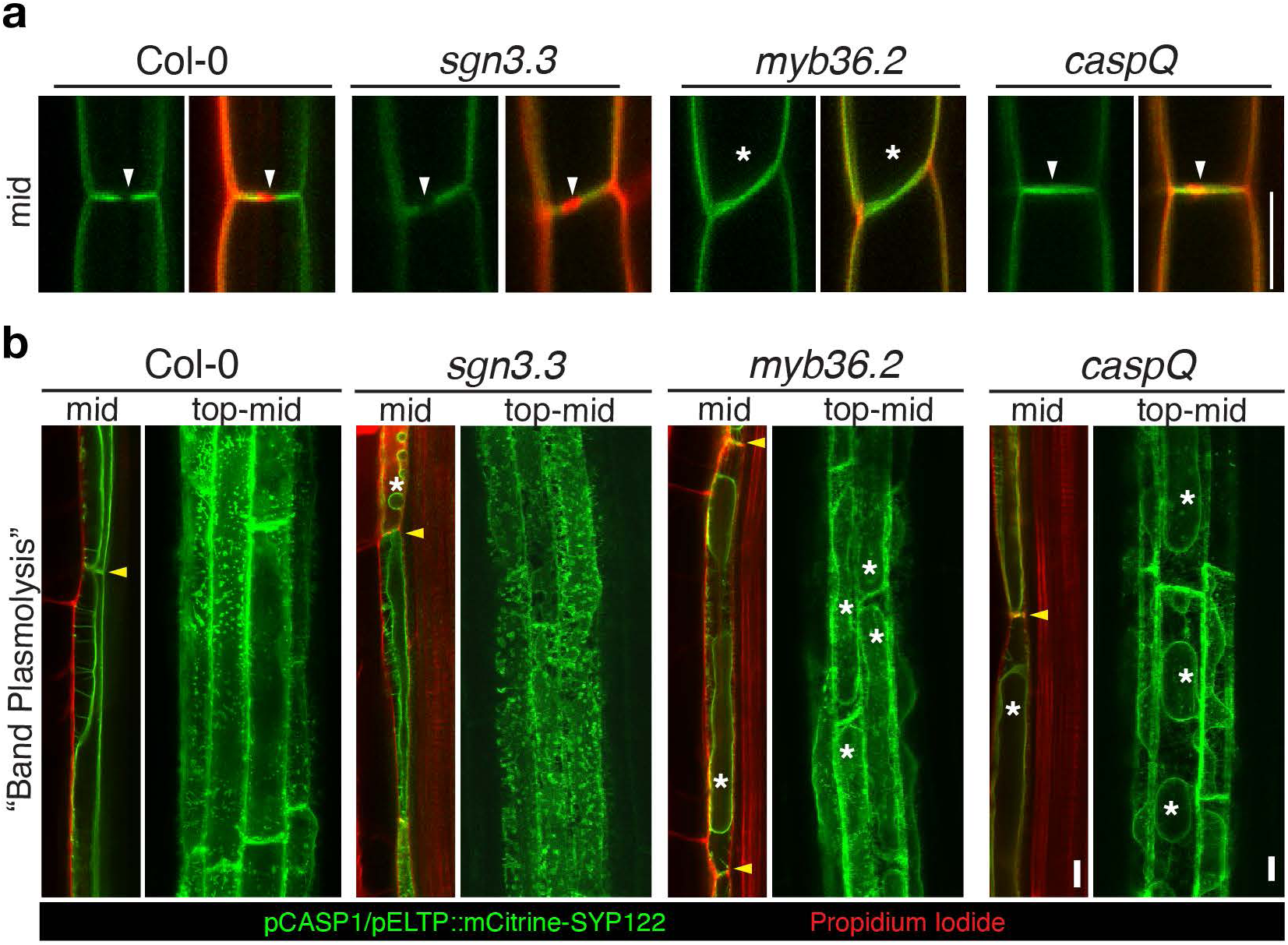
CSD exclusion zone and plasma membrane wall attachment after plasmolysis in Col-0, sgn3, myb36 and caspQ. **a.** Plasma-membrane marker mCitrine-SYP122 is excluded from CSD labelled by the Propidium Iodide-stained CS in Col-0 and *sgn3* (white arrows) but is freely diffused in transversal membranes of *myb36* and of *caspQ*, with difference that *myb36* lacks a CS (asterisk) but *caspQ* still forms a CS (white arrow). **b.** Plasma membrane marker mCitrine-SYP122 and PI-stained cell-walls after plasmolysis in seedlings mounted in 0.8M Mannitol solution. For Col-0 in mid-view band-plasmolysis can be observed (yellow arrow), and but no obvious detachment appears in top-mid view. In *sng3* mid-view protoplasts (*) but in top-mid view only mild detachments. *caspQ* and *myb36* display very similar behaviour, with clear protoplasts (*) in either mid o trop-mid views.

**Supplementary Figure 5.1:**
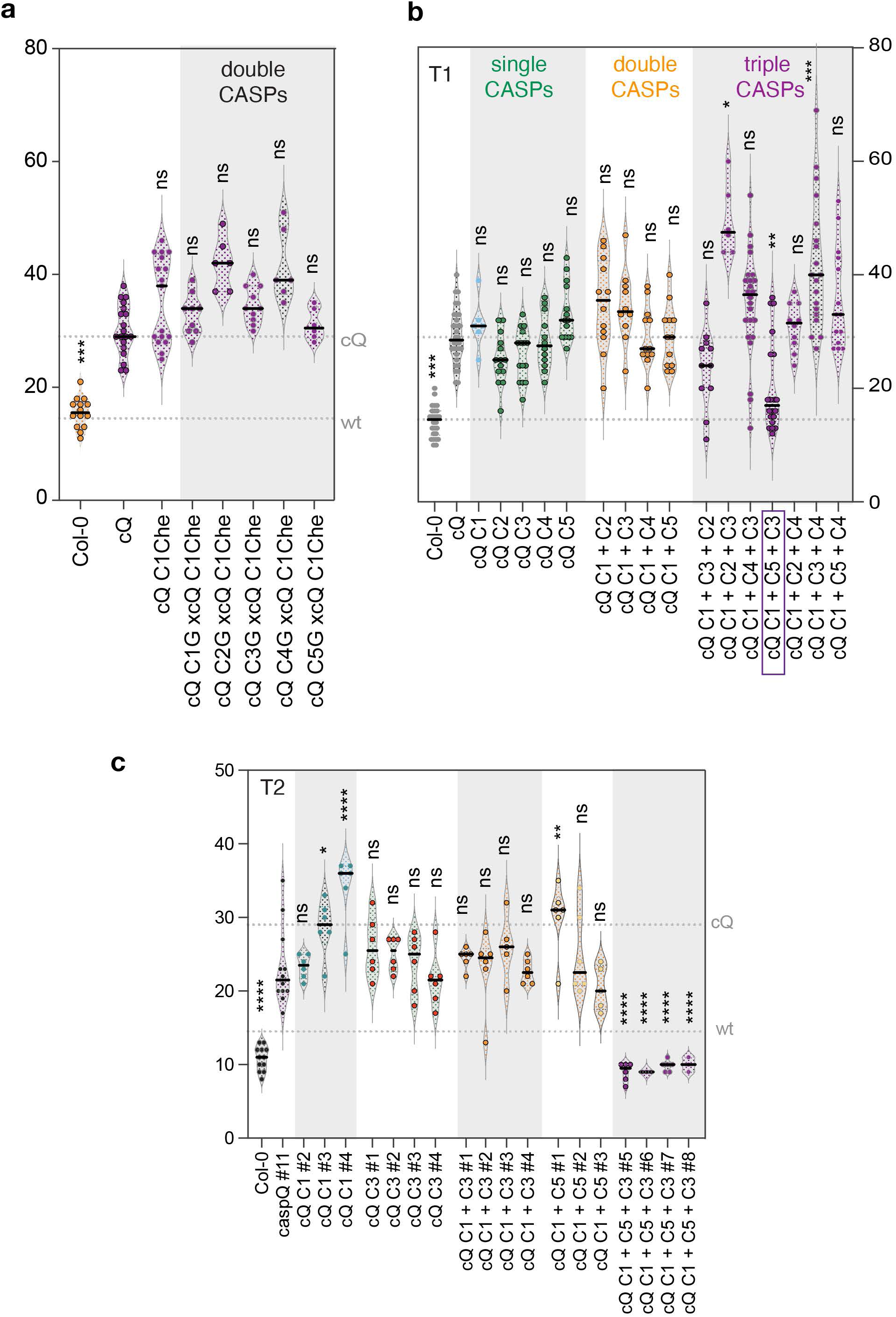
Multiple CASP combinations reveals caspQ complementation requires at least three CASP genes. **a-b.** Propidium iodide uptake assay in *caspQ* complementation lines **(a):** double CASP combinations obtained by crossing caspQ + pCASP1:CASP1-mCherry with caspQ + pCASPn:CASPn-GFP. **(b-c):** single, double and triple CASP1-5 complementation of caspQ obtained by combinatorial gateway constructs with one, two or three CASPs in T1, n=12-15 individuals (b), and in T2 for a selection of constructs in individuals selected for presence of transgene, n=6-8 individuals (c). Statistical differences by ANOVA, Dunnet’s test comparisons to *caspQ* (p<0.05). Highlighted with purple box is the best complementing CASP combination: 1 + 5 + 3, consistent among independent T1 individuals and reproducible in T2.

**Supplementary Figure 6.1:**
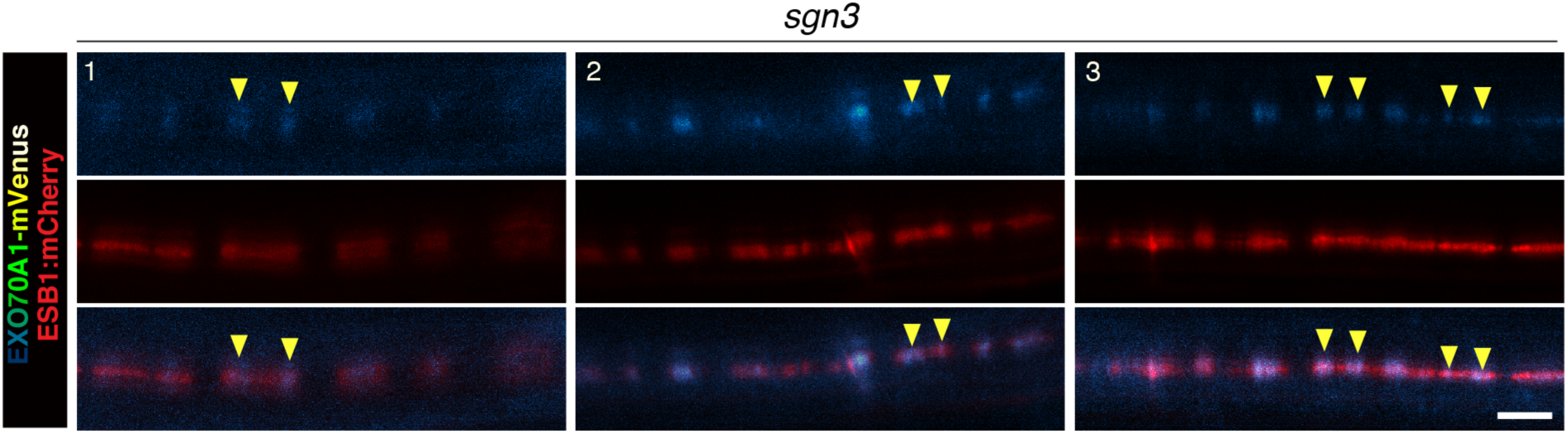
EXO70A1-mVenus is partially displaced in sgn3. **a.** Three representative seedlings for the localization of EXO70A1-mVenus and ESB1-mCherry in CS surfaces of *sgn3* cell number 9. Scale bar 5 µm.

**Supplementary Figure 7.1:**
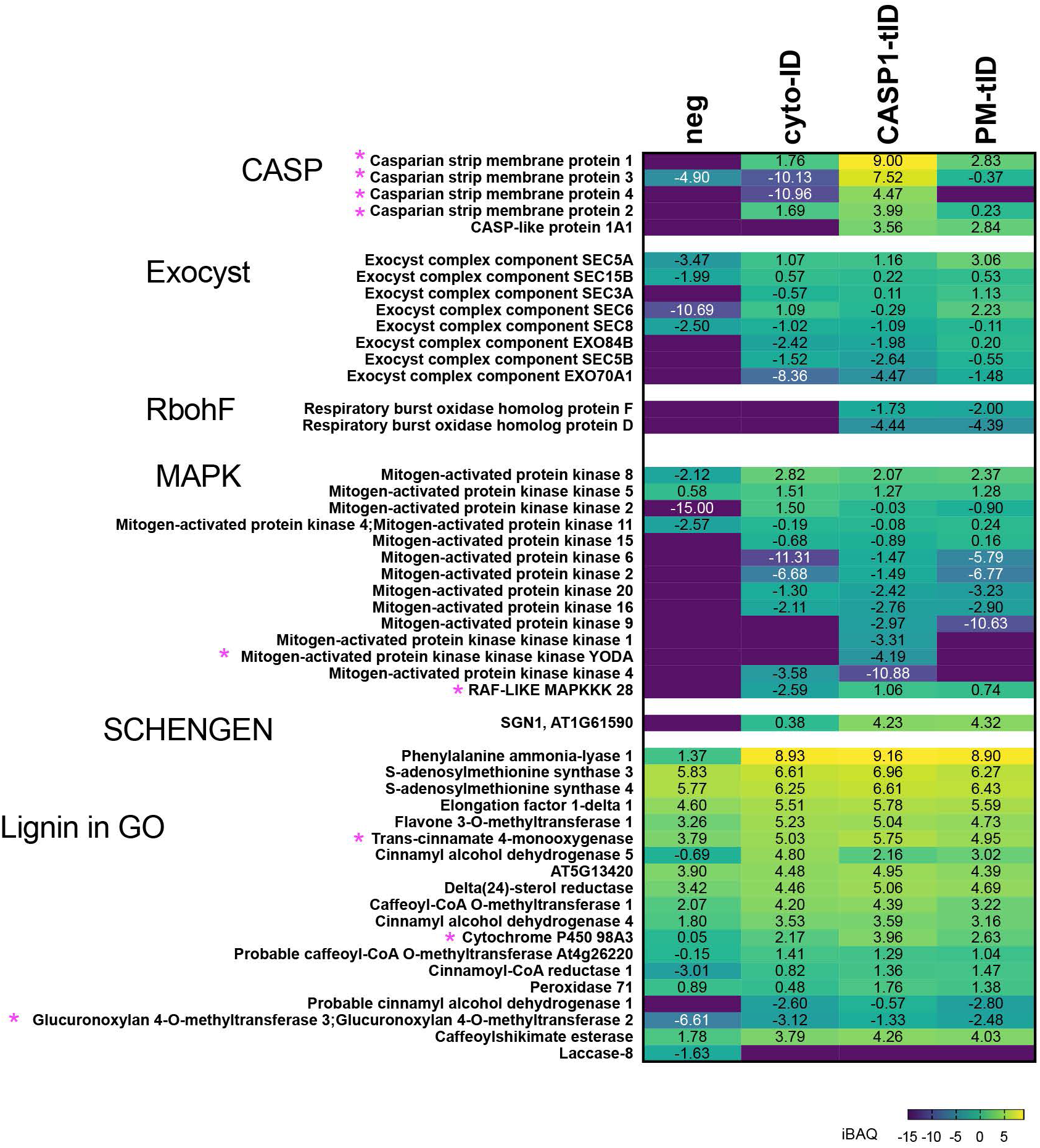
Profile of CS associated proteins in Endodermal turboID proximity labelling experiment. **a.** Detected members, abundances in TurboID samples (log2 iBAQ) and their significant enrichment (p<0.05, paired t-test Bonferroni adjusted *q*-value) in CASP1-tID sample (asterisk), from protein families or GO-terms expected to be involved in CS formation. *CASP-Like family*: 5 out of 39 members were identified in the experiment, four of which (CASP1 to CASP4) significantly enriched in CASP1-tID. *Exocyst*: one member of each subunit of exocyst complex was detected, with exceptions of subunit …. For which no member was identified; *RBOH* (Respiratory Oxidative Burst Homologue): two isoforms involved in CS (RBOHF) and CS-surveillance endodermal lignification (RBOHD) (Fujita et al 2020); *Mitogen-Activated protein kinases:* detected members from MAPKKK, MAPKK and MPK families; *SCHENGEN* pathway, i.e. SGN1 and SGN3, only SGN1 was identified; and Lignin in GO-term. Neg (negative control, pCASP1::CASP1-GFP; cytoID, pCASP1::GFP-turboID; CASP1tID, pCASP1::CASP1-GFP-turboID, and PM-tID, pCASP1::GFP-turboID-SYP122).

**Supplementary Figure 7.2:**
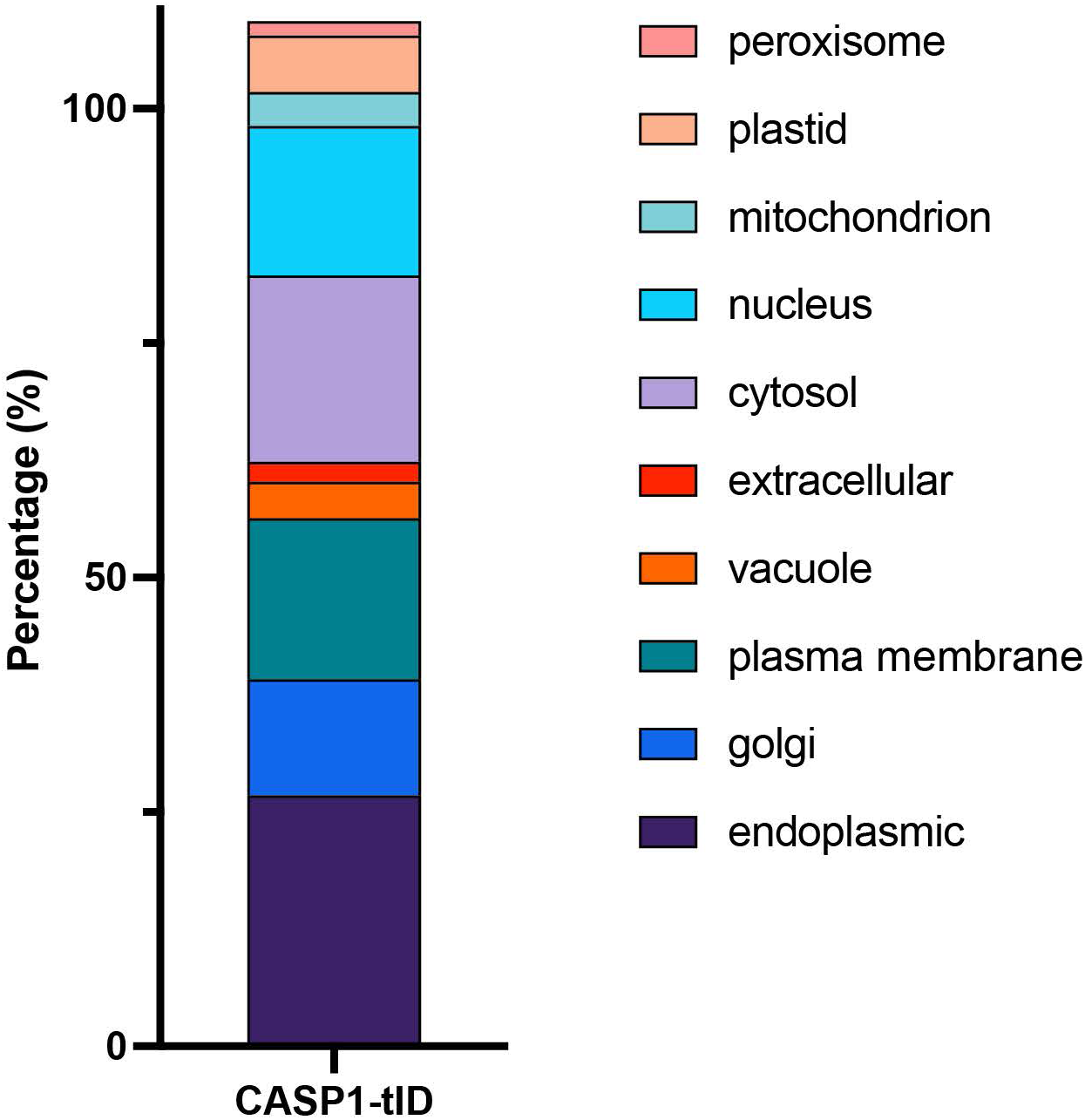
Predicted sub-cellular localization of CASP1-proximal proteins. **a.** Relative (%) sub-cellular localizations of 332 CASP1-tID enriched proteins as predicted by SUBA4, consensus method. Note many proteins have predicted dual/multiple localisations, thus sum of % is above 100%.

**Supplementary Figure 7.3:**
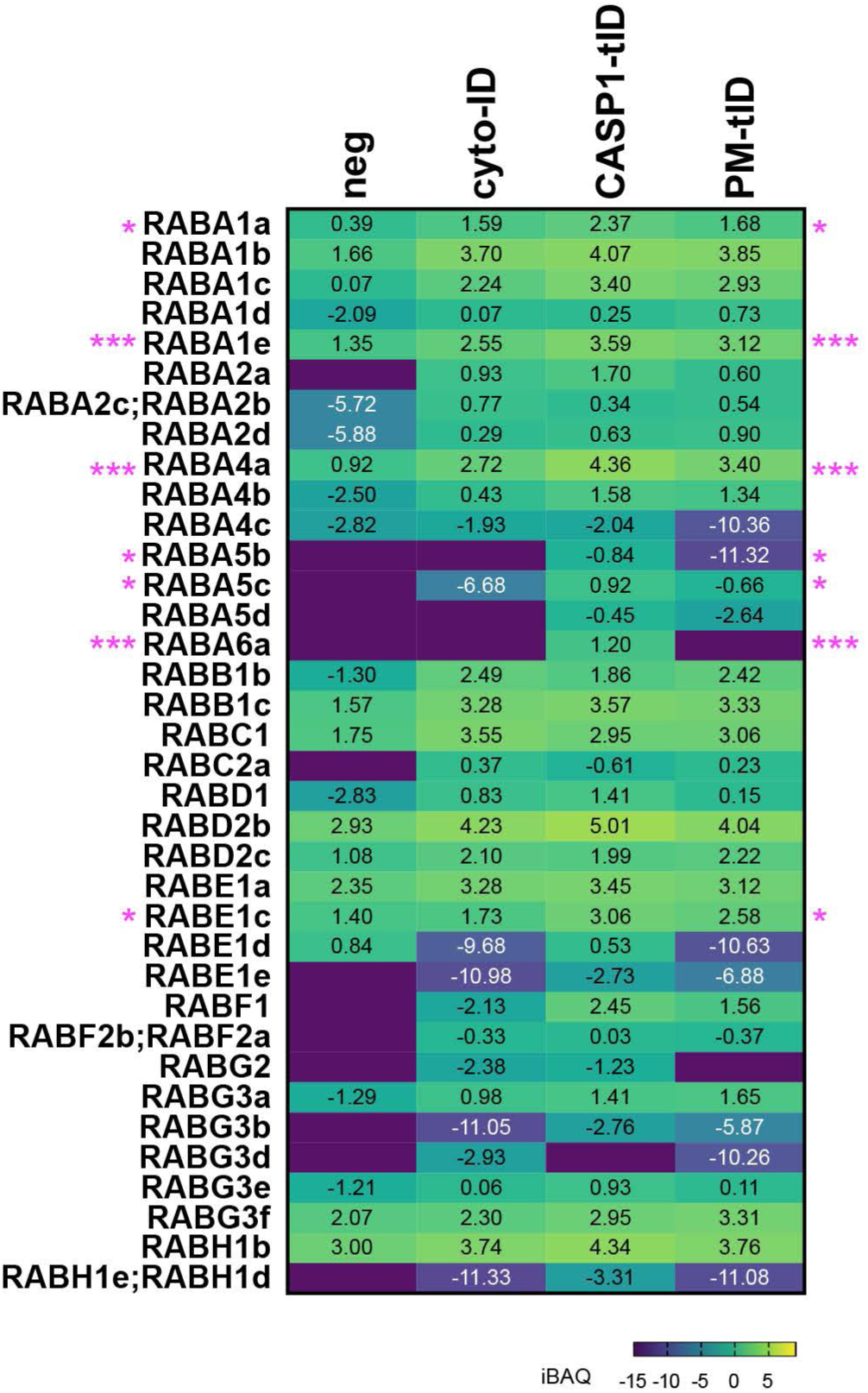
Profile of Rab-GTPase family in Endodermal turboID proximity labelling experiment. **a.** Detected members, protein abundances in TurboID samples (log2 iBAQ) and significant enrichment (p<0.05) in CASP1-tID sample (asterisk) from Rab-GTPase family.

